# Dual Regulatory Role of Nuclear FNBP4 in Actin Binding and Formin FMN1 Inhibition

**DOI:** 10.64898/2026.01.05.697755

**Authors:** Shubham Das, Saikat Das, Sankar Maiti

## Abstract

Nuclear actin dynamics are increasingly recognized as central to genome regulation, yet the identity and function of dedicated nuclear actin regulators remain elusive. Here, we uncover formin-binding protein 4 (FNBP4) as a previously uncharacterized nuclear actin-binding protein that integrates dual regulatory functions. We identify a conserved basic KRRK motif within its C-terminal that directly interacts actin and coincides with a non-canonical nuclear localization signal. In parallel, FNBP4 interacts with the formin FMN1 via its N-terminal WW domains, potently suppressing FMN1-mediated actin nucleation. Strikingly, full-length FNBP4, which unites both the FMN1- and actin-binding modules, displays synergistic inhibition far exceeding that of truncated fragments. Furthermore, confocal imaging confirms nuclear colocalization of FNBP4 with actin, highlighting its role in shaping nuclear actin architecture. Our findings position FNBP4 as the first example of a nuclear-resident formin inhibitor that simultaneously binds actin and a formin. This dual functionality establishes a new regulatory paradigm in which a single protein integrates direct actin interaction with formin inhibition to fine-tune nuclear actin organization.

## 1. Introduction

Dynamic remodeling of the actin cytoskeleton is essential for various cellular processes, including cell shape regulation, migration, and intracellular trafficking (Blanchoin et al., 2014; Pollard and Cooper, 2009; Reisler and Egelman, 2007). While the cytoplasmic roles of actin are well-established, increasing evidence highlights the presence and functional significance of actin within the nucleus, where it contributes to chromatin remodeling, transcriptional regulation, and nuclear architecture (“Dynamizing nuclear actin filaments,” 2019; Ulferts et al., 2024; Wollscheid and Ulrich, 2023). Although actin lacks a classical nuclear localization signal (NLS), it actively shuttles between the nucleus and cytoplasm through a regulated transport system, reflecting the existence of a shared actin pool between these compartments (Kristó et al., 2016; Vartiainen, 2008). Notably, certain actin-binding proteins facilitate this nucleocytoplasmic trafficking. For example, cofilin, in complex with importin-9, mediates the nuclear import of actin, while profilin, in association with exportin-6, facilitates its export back to the cytoplasm (Dopie et al., 2012; Stüven et al., 2003). Several actin-binding proteins shuttle between the cytoplasm and nucleus to maintain nuclear actin dynamics, particularly during key nuclear processes such as DNA damage response, maintenance of genome stability, and transcriptional regulation (Kristó et al., 2016). For example, Junction-mediating and regulatory protein (JMY), which localizes to the cytoplasm under normal conditions to promote actin filament assembly and facilitate cell migration, translocates to the nucleus in response to DNA damage, an actin-dependent process (Zuchero et al., 2012). Another group of proteins actin nucleators like mDia2 continuously shuttles between the nucleus and cytoplasm (Miki et al., 2009). FMN2 accumulated in the nucleus to promote actin polymerization during the DNA damage response as well as under hypoxic conditions (Belin et al., 2015; Yamada et al., 2013). However, the molecular mechanisms that coordinate actin dynamics within the nuclear compartment remain incompletely understood. In particular, the contribution of exclusively nuclear-localized proteins to the regulation of nuclear actin architecture is an emerging area that remains insufficiently characterized.

Several ABPs not only modulate actin dynamics but also interact directly with formins, thereby exerting dual regulatory roles. For instance, Drebrin, primarily expressed in neurons, contains an actin-binding domain spanning residues 233 to 317, which contributes to the organization of the dendritic actin network, while its intrinsically disordered C-terminal region interacts with the FH2 domain and the C-terminal tail of the formin protein mDia2 (Ginosyan et al., 2019; Grintsevich et al., 2010; Mikati et al., 2013; Srapyan et al., 2023; Worth et al., 2013). Another example, Flightless I (Flii), classified within the broader group of actin-regulatory proteins related to gelsolin, is characterized by leucine-rich repeats (LRRs) at its N-terminal that mediate interaction with actin (Liu and Yin, 1998; Mohammad et al., 2012). In addition to its actin-binding capability, Flii enhances actin assembly by directly interacting with the Diaphanous Autoregulatory Domain (DAD) of Diaphanous-related formins (DRFs), such as Daam1 and mDia1 (Higashi et al., 2010). Furthermore, IQGAP1, a multifunctional scaffold protein, interacts with Diaphanous-related formins via the Diaphanous Inhibitory Domain (DID) of DIAPH1 at the leading edge of migrating cells, and also facilitates F-actin cross-linking into a mesh-like network, and has recently been shown to cap the barbed ends of F-actin (Chen et al., 2020; Liu et al., 2016; Mateer et al., 2004; Pimm et al., 2024). Profilin, a well-characterized actin monomer-binding protein, enhances formin-mediated filament elongation by delivering profilin–actin complexes to the growing filament barbed ends by interacting with the FH1 domain of formins (Paul and Pollard, 2008; Perelroizen et al., 1994; Pring et al., 2003; Schutt et al., 2022). Anillin, a multifunctional scaffolding protein involved in various cytoskeletal structures, binds to F-actin to promote filament bundling and interacts with the Diaphanous Inhibitory Domain (DID) of DIAPH3 to facilitate its activation (Chen et al., 2017; Field and Alberts, 1995; Piekny and Glotzer, 2008). Formin-binding proteins, including Drebrin, IQGAP1, Flii, Anillin, and Profilin, act as molecular bridges linking formin activity to actin dynamics. However, the actin-binding activities of other formin-binding proteins remain poorly understood, leaving an important question open. Furthermore, identifying regulators that not only modulate formin function but also directly influence nuclear actin dynamics presents an intriguing avenue for understanding the mechanisms governing actin regulation within the nucleus.

Formin-binding protein 4 (FNBP4) is a WW domain-containing protein that has been implicated in a range of physiological processes. Notably, its expression is transcriptionally upregulated in response to cellular stress triggered by γ-irradiation, suggesting a role in stress-responsive signaling pathways (Depraetere and Golstein, 1999). Emerging evidence also highlights FNBP4 as a potential biomarker linked to cuproptosis, where it has been shown to promote hepatocellular carcinoma (HCC) cell proliferation and metastasis, further underscoring its relevance in cancer biology (Zheng et al., 2023). Furthermore, FNBP4 has been associated with Microphthalmia with Limb Anomalies (MLA), also known as Waardenburg anophthalmia syndrome or ophthalmoacromelic syndrome (Kondo et al., 2013). This rare autosomal recessive disorder is characterized by developmental defects affecting the eyes and limbs in both humans and mice, suggesting a critical role for FNBP4 in embryonic development and tissue patterning (Kondo et al., 2013). Our previous studies established a direct interaction between the N-terminal WW1 domain of FNBP4 and the FH1 domain as well as the FH1-linker region of FMN1, highlighting their specific binding (Das and Maiti, 2024; Das et al., 2025). Furthermore, we demonstrated that FNBP4 as a nuclear protein and acts as a stationary inhibitor of FMN1, suppressing FMN1-driven actin cytoskeleton dynamics (Das et al., 2025). However, whether FNBP4 directly interacts with actin and functions in a dual regulatory capacity, akin to other actin-binding adaptor proteins, remained unclear.

In this study, we systematically dissected the interaction profile of FNBP4 with actin, using a series of truncation constructs, co-sedimentation assays, and molecular docking analyses. Our results reveal that FNBP4 as an actin-binding protein, and mutational analyses identified the C-terminal region of FNBP4 as critical for the direct actin-binding site, with key residues KRRK (1009-1012 amino acids) mediating the interaction. Furthermore, extreme C-terminal region harbors a basic KRRK motif (1009-1012 amino acids) that directly binds F-actin and inhibits actin polymerization in vitro. Confocal imaging confirmed the nuclear colocalization of FNBP4 with actin, suggesting a previously unrecognized role for FNBP4 in modulating nuclear actin dynamics. Furthermore When the FMN1-interacting region and the actin-binding region are present together, the inhibitory activity of full-length FNBP4 is more potent than that of the N-ter WW1-WW2 FNBP4 fragment. Together, these findings identify FNBP4 as a dual regulator of actin cytoskeleton remodeling through its interaction with both FMN1 and uncover a novel actin-binding motif within its C-terminal with potential nuclear functions.

## 2. Result

### 2.1. Differential function of FNBP4 termini: N-terminal interacts with FMN1 and C-terminal binds actin

Our previous biochemical investigations established that the WW1 domain (resides in N-terminal) of FNBP4 interact with the FH1 domain and the interdomain connecter region between FH1 and FH2 domain of FMN1 (Das et al., 2025). This interaction appears to regulate FMN1-mediated actin cytoskeleton dynamics. Moreover, these studies also revealed that the WW1-WW2 FNBP4 (214-629 amino acids) fragment does not exhibit any direct interaction with actin, suggesting that the WW domains may specifically modulate FMN1 function without engaging actin independently. Interestingly, several known actin regulators such as Debrin, Profilin, Ena/VASP, Spire1/2, Tropomyosin, Cap1/2, Twinfilin and IQGAP1 can bind both formins and actin, thereby contributing to the dynamic remodeling of the actin cytoskeleton through dual interactions. These observations prompted us to investigate whether FNBP4 could similarly function in a dual regulatory capacity, by modulating FMN1 activity and also through direct interaction with actin. To systematically examine the potential actin-binding properties of FNBP4, we generated two truncation constructs covering the N-terminal and C-terminal regions of the full-length FNBP4 protein (Fig. 1A). The first construct, termed N-ter FNBP4, includes amino acid residues 1-647, encompassing the tandem WW domains. The second construct, C-ter FNBP4, comprises residues 647-1017, representing the C-terminal half of FNBP4 that lacks WW domains but contain other two putative monopartite nuclear localization signals (Fig. 1A). Then both N-ter FNBP4 and C-ter FNBP4 constructs were purified from bacterial system as 6×His-tagged fusion protein (Fig. S 1B).

**Figure 1.**
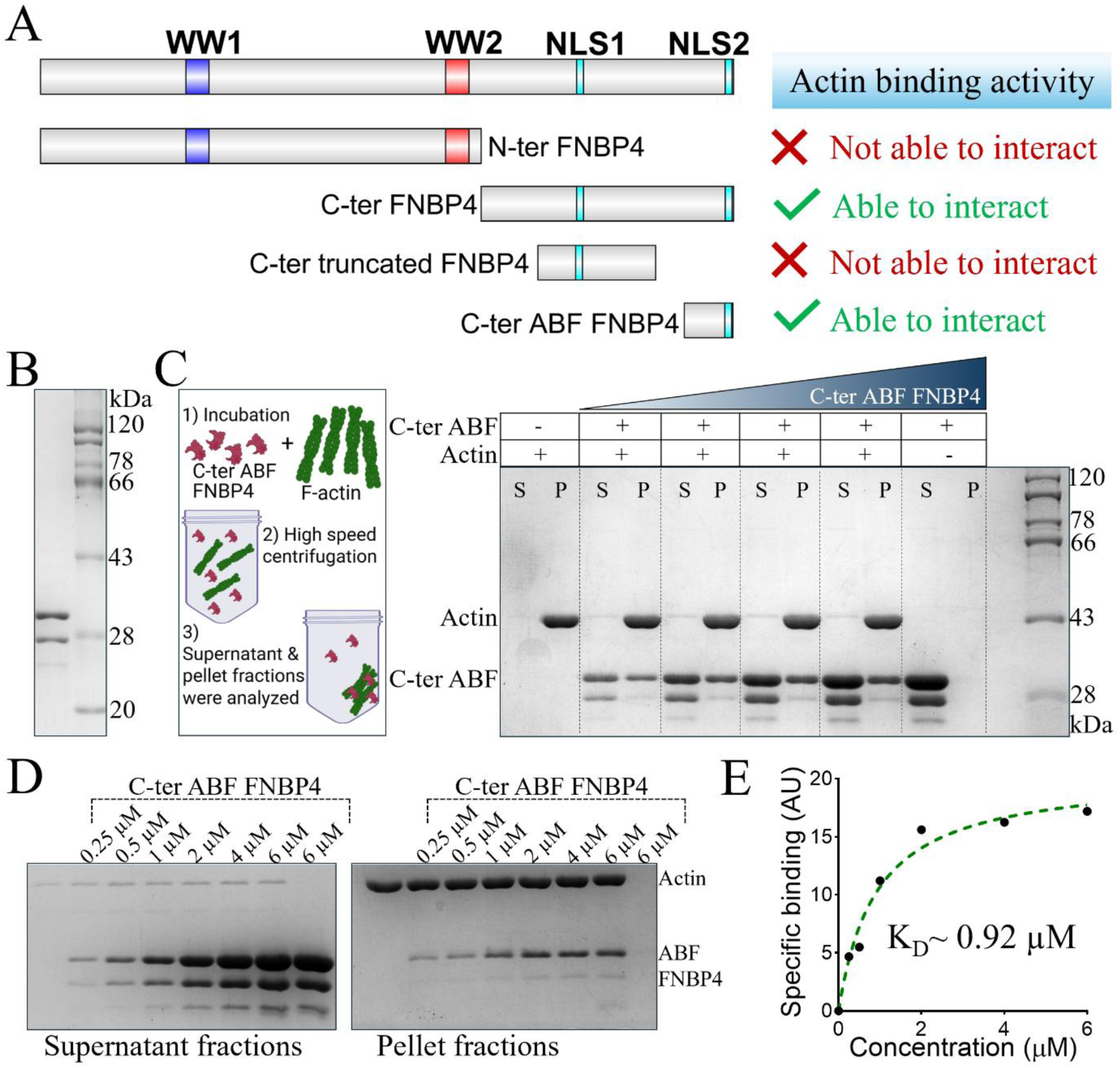
C-terminal ABF region of FNBP4 exhibits actin-binding activity. (A) Schematic representation of FNBP4 domain organization and the design of truncation constructs. Constructs include N-terminal FNBP4 (1-647 amino acids), C-terminal FNBP4 (647-1017 amino acids), and truncated C-terminal FNBP4 (732-904 amino acids) and C-terminal ABF FNBP4 (946-1017 amino acids). The corresponding actin-binding activities for each constructs are indicated in the figure. (B) Coomassie-stained SDS-PAGE analysis of purified GST-tagged C-terminal ABF FNBP4. (C) F-actin co-sedimentation assay. A schematic of the co-sedimentation assay is included to illustrate the experimental setup. Polymerized actin (5 µM) was incubated in the absence or presence of the C-terminal ABF FNBP4 fragment (946-1017 amino acids) for 15 min at room temperature. Samples were then subjected to ultracentrifugation, and the supernatant (S) and pellet (P) fractions were analyzed by 10% SDS-PAGE and visualized with Coomassie staining. The presence of C-terminal ABF FNBP4 in the pellet with F-actin indicates a specific interaction. (D) Quantitative analysis of the binding interaction between the C-terminal ABF FNBP4 fragment and F-actin. Increasing concentrations of C-terminal ABF FNBP4 (0.25 µM, 0.5 µM, 1 µM, 2 µM, 4 µM and 6 µM) were incubated with 2.5 µM F-actin, followed by ultracentrifugation. Supernatant (S) and pellet (P) fractions were analyzed separately by 10% SDS-PAGE. (E) Graph shows the pellet fraction, representing bound protein, quantified by band intensity (arbitrary units, AU) on the Y-axis and plotted against the C-terminal ABF FNBP4 concentration (µM) on the X-axis. Data were fitted to a one-site binding model, yielding a dissociation constant (K_D_) approximately 0.92 µM.

These purified truncation constructs were then subjected to further in vitro assays to assess their actin-binding capabilities. We performed F-actin co-sedimentation assays using the purified N-terminal and C-terminal truncation constructs (Fig. S 1C and D). In this assay, pre-polymerized F-actin was incubated with each FNBP4 fragment, followed by ultracentrifugation to separate the supernatant (unbound proteins) and pellet (F-actin and associated proteins) fractions. The results revealed that the N-terminal FNBP4 fragment remained in the supernatant fraction, indicating that it does not interact with F-actin under the assay conditions (Fig. S 1C). In contrast, the C-terminal FNBP4 was detected in the pellet fraction along with F-actin, demonstrating a clear interaction between the C-terminal region of FNBP4 and F-actin (Fig. S 1D). These findings suggest that actin-binding activity is localized to the C-terminal region of FNBP4, whereas the N-terminal region lacks such capability but instead mediates interactions with FMN1.

### 2.2. Mapping the actin-binding region within the C-terminal of FNBP4

To further delineate the specific region within the C-terminal of FNBP4 responsible for actin binding, the C-terminal fragment (647-1017 amino acids) was subdivided into two smaller fragments: C-terminal truncated FNBP4 (732-904 amino acids) and C-terminal ABF FNBP4 (946-1017 amino acids) (Fig. 1A). These constructs were designed to test whether the actin-binding functionality resides within a specific subregion of the C-terminal. C-terminal truncated FNBP4 fragment was expressed and purified as a 6×His-tagged fusion protein (Fig. S 1E) and C-terminal ABF FNBP4 fragment was expressed as a GST-tagged fusion protein, and purified with an approximate molecular weight of 34 kDa (Fig. 1B). Here we find that this fragment is prone to degradation during purification, with two additional degradation bands appearing below the main band (Fig. 1B). To assess their actin-binding abilities, F-actin co-sedimentation assays were performed with both fragments. The results showed that the C-terminal truncated FNBP4 remained in the supernatant fraction after ultracentrifugation, indicating it does not interact with F-actin (Fig. S 1F). In contrast, the C-terminal ABF FNBP4 fragment was found in the pellet along with F-actin, confirming its ability to bind to actin filaments (Fig. 1C). Moreover, GST alone was included as a negative control and remained entirely in the supernatant (Fig. S 1G), demonstrating that the observed actin interaction is specific to the FNBP4 and not due to the GST tag.

To quantify the binding affinity, a binding assay was conducted using increasing concentrations of the C-terminal ABF FNBP4 fragment with a constant concentration of F-actin (2.5 µM) (Fig. 1D). The band intensities of pellet fractions were analyzed using ImageJ (Fiji). The bound protein fraction was plotted against the total protein, and the binding curve was fitted using a one-site specific binding model. From this analysis, the dissociation constant (K_D_) for the interaction between the C-terminal ABF FNBP4 fragment and F-actin was determined to be approximately 0.92 µM, indicating a moderate binding affinity (Fig. 1E). These results clearly identify the 946-1017 region of FNBP4 as a functional actin-binding region, suggesting that this discrete region within the C-terminal portion is responsible for mediating direct interaction with actin filaments.

### 2.3. Molecular Docking Predicts Actin-Binding Interface Within the C-terminal ABF Region of FNBP4

To gain structural insight into the interaction between C-ter ABF FNBP4 and actin, we conducted a molecular docking analysis using the predicted 3D structure of the C-terminal ABF region of FNBP4 (946-1017 amino acids) (Fig. S 2A). The structure was modeled using AlphaFold, which provided a high-confidence prediction for the fragment (Fig. S 2B). This model was then subjected to protein-protein docking using the HADDOCK web server, with actin as the binding partner. The docking results revealed several potential binding poses, among which the top-ranked cluster displayed a stable and plausible interaction interface between the FNBP4 C-terminal fragment and actin (Fig. 2A). To identify key residues involved in the interaction, we analyzed the interface using LigPlot+. This analysis highlighted a short stretch of basic residues, Lys1009, Arg1010, Arg1011, and Lys1012 (KRRK motif), as central to the actin-binding interface (Fig. 2A and S 2C). Further, to evaluate the stability and conformational dynamics of the C-terminal ABF FNBP4-actin complex, we performed a 50 ns molecular dynamics (MD) simulation using GROMACS. The root mean square deviation (RMSD) analysis indicated that the complex stabilized at ∼ 0.808 nm toward the end of the simulation (Fig. 2B). The radius of gyration (Rg), which reflects the overall compactness of the system, was calculated to be 2.503 nm, with component values of Rgx = 1.706 nm, Rgy = 2.310 nm, and Rgz = 2.069 nm (Fig. 2C). Collectively, these results suggest that the C-terminal ABF FNBP4 forms a stable and compact complex with actin, thereby reinforcing its structural persistence. These findings provide computational evidence that the KRRK motif in the extreme C-terminal region of FNBP4 may mediate direct interaction with actin, supporting our biochemical observations of actin binding by this fragment.

**Figure 2.**
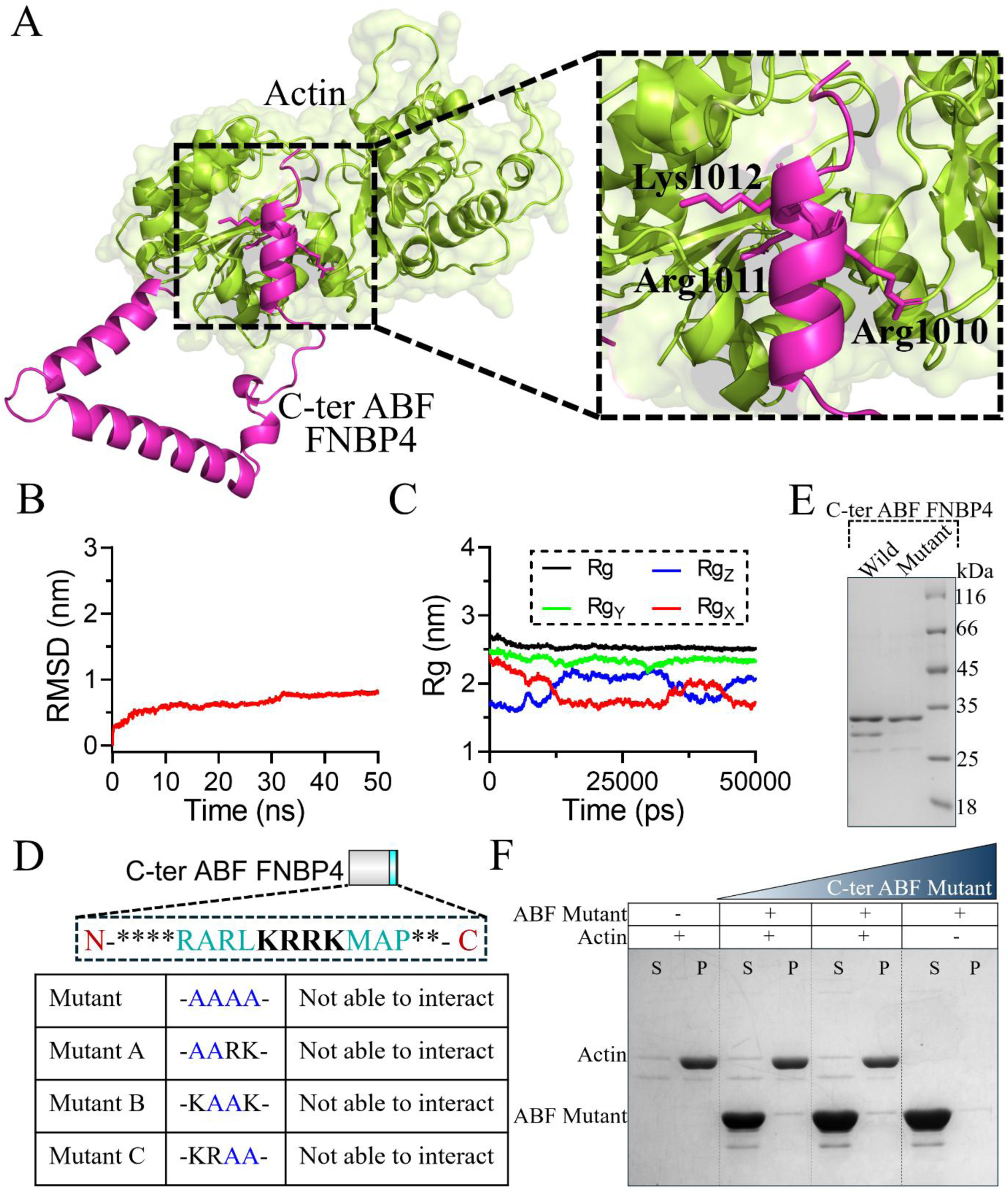
KRRK motif is responsible for interacting with actin. (A) The top-scoring docked model of actin (lime green) bound to the C-terminal ABF FNBP4 (light magenta) is shown. The inset provides a close-up of the binding interface, highlighting key residues predicted to mediate the interaction and contribute to complex formation. (B) Backbone RMSD of the actin–C-terminal ABF FNBP4 complex over a 50-ns molecular dynamics simulation, showing structural stability throughout the trajectory. (C) Radius of gyration (Rg) analysis of the actinand C-terminal ABF FNBP4 complex during a 50 ns molecular dynamics simulation. Overall Rg is shown in black, while Rg values along the x, y, and z axes are represented in red (Rg_X_), green (Rg_Y_), and blue (Rg_Z_), respectively. The consistent Rg values indicate that the complex maintains a stable and compact conformation throughout the simulation. (D) C-terminal ABF FNBP4 mutants used for actin-binding analysis. The wild-type C-terminal ABF FNBP4 and four mutants were cloned into the pGEX-4T3 vector. Mutants include: C-terminal ABF Mutant FNBP4 (KRRK-to-AAAA; 1009-1012 amino acids), MutantA (KR-to-AA; 1009-1010 amino acids), MutantB (RR-to-AA; 1010-1011 amino acids), and and MutantC (RK-to-AA; 1011-1012 amino acids). The corresponding actin-binding activities for each mutant are indicated in the figure. (E) Coomassie-stained SDS-PAGE analysis of purified GST-tagged C-terminal ABF FNBP4, C-terminal ABF Mutant FNBP4. (F) F-actin co-sedimentation assay: Polymerized actin (5 µM) was incubated with or without C-terminal ABF Mutant FNBP4 (KRRK-to-AAAA; 1009-1012 amino acids) for 15 minutes at room temperature. Samples were then subjected to ultracentrifugation, and the resulting supernatant (S) and pellet (P) fractions were analyzed by 10% SDS-PAGE, followed by Coomassie staining for visualization.

To validate this prediction, a mutant construct of the C-terminal ABF FNBP4 fragment was designed in which the KRRK motif (1009-1012 amino acids) was substituted with alanines (KRRK-to-AAAA), effectively neutralizing the positive charge of this region (Fig. 2D). The mutant fragment was expressed and purified as a GST-tagged fusion protein with a similar molecular weight like wild type C-terminal ABF FNBP4 (Fig. 2E). Subsequently, the effect of the KRRK-to-AAAA mutation on actin binding, we performed an F-actin co-sedimentation assay under identical conditions to those used for the wild-type fragment (Fig. 2F). The results showed that, unlike the wild-type C-terminal ABF FNBP4 fragment, which co-sedimented with F-actin and was found predominantly in the pellet fraction, the mutant fragment was detected exclusively in the supernatant (Fig. 2F). This indicated that the mutant lost its ability to bind actin filaments. These results clearly demonstrate that the KRRK motif (1009-1012 amino acids) within the extreme C-terminal region of FNBP4 is essential for actin binding. Loss of interaction upon mutation of this motif highlights its critical role in mediating the direct association between FNBP4 and actin.

To further delineate the importance of individual residues within the KRRK motif (1009-1012 amino acids) of the FNBP4 C-terminal ABF region in mediating actin binding, we generated a series of double-point mutants by replacing specific pairs of residues with alanines. Three constructs were designed: MutantA (KR-to-AA; 1009-1010 amino acids), MutantB (RR-to-AA; 1010-1011 amino acids), and MutantC (RK-to-AA; 1011-1012 amino acids) (Fig. 2D and S 3A). Each mutant was expressed and purified as a GST-tagged protein and subjected to F-actin co-sedimentation assays under identical conditions as the wild-type ABF fragment (Fig. S 3C, D and E). The results showed that none of the three mutant fragments were able to co-sediment with F-actin. All mutant proteins were exclusively detected in the supernatant fraction, indicating a complete loss of actin-binding capability (Fig. S 3C, D and E). These observations suggest that the integrity of the full KRRK motif is essential for FNBP4’s interaction with actin, and disruption of any part of this basic stretch is sufficient to abolish binding. This further supports the conclusion that the KRRK motif (1009-1012 amino acids) within the extreme C-terminal region of FNBP4 constitutes a critical actin-binding determinant. These residues are located near the extreme C-terminal end of FNBP4, overlapping with the previously predicted NLS2 (RARL**KRRK**MA; 1005-1015 amino acids), which intriguingly lacks nuclear localization function in isolation.

### 2.4. C-teriminal ABF FNBP4 Fragment Interacts with G-actin and Inhibits Actin Polymerization

We evaluated the functional relevance of C-ter ABF FNBP4’s actin-binding capacity by performing end-point actin polymerization assays using Total Internal Reflection Fluorescence (TIRF) microscopy. G-actin was incubated with either the wild-type C-ter ABF FNBP4 fragment or the C-ter ABF mutant FNBP4 fragment (KRRK-to-AAAA) at a final concentration of 4 µM (Fig. 3A and B). Actin polymerization was monitored by imaging the filaments at 5 and 10 minutes under TIRF microscopy. The results showed that the wild-type C-ter ABF FNBP4 fragment significantly inhibited actin polymerization, as evidenced by a marked reduction in the number of filaments observed at both time points compared to the actin-alone control (Fig. 3A). In contrast, the C-ter ABF mutant FNBP4 fragment had no observable effect on actin polymerization; the number of filaments remained comparable to that of the actin control (Fig. 3A). To quantify the results, the number of actin filaments was manually counted from the TIRF images under each condition. The filament counts were plotted as a bar graph showing filament number versus the respective experimental conditions (Actin only, Actin + C-ter ABF FNBP4, Actin + Mutant C-ter ABF FNBP4) (Fig. 3B). The data clearly demonstrated that the wild-type C-ter ABF FNBP4 fragment suppresses actin filament formation, while the mutant fragment, which lacks actin-binding due to the mutated KRRK motif, does not exhibit this inhibitory effect.

**Figure 3.**
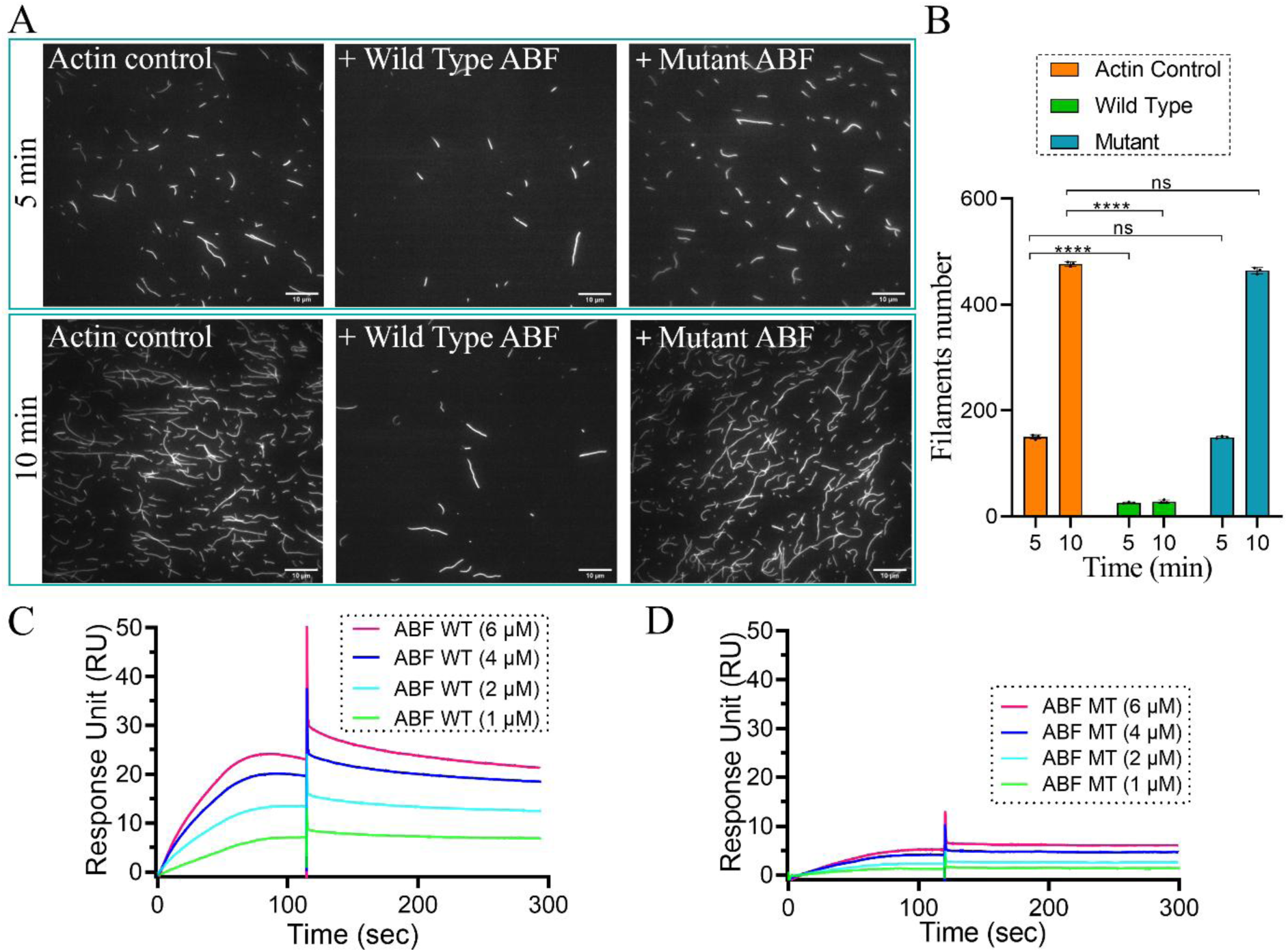
The C-terminal ABF region of FNBP4 interacts with monomeric G-actin and inhibits actin polymerization via its KRRK motif. **(A)** Time-lapse TIRF microscopy images of actin filament polymerization. G-actin (3 µM) was polymerized in the absence or presence of 4 µM wild-type C-ter ABF FNBP4 or 4 µM C-terminal ABF Mutant FNBP4 (KRRK-to-AAAA; 1009-1012 amino acids). Actin filaments were stained with Alexa Fluor 488–phalloidin and imaged at 5 and 10 min from separate experiments. Scale bar: 10 µm. **(B)** Quantification of filament numbers at the endpoint from three independent fields. Wild-type C-ter ABF FNBP4 significantly reduced filament formation, whereas the mutant fragment did not affect polymerization. Error bars represent the standard deviation. Statistical significance was determined using an unpaired two-tailed Student’s *t*-test in GraphPad Prism 8. Significance is indicated as *P ≤ 0.05, **P ≤ 0.01, ***P ≤ 0.001, ****P ≤ 0.0001, and ns (not significant). (C) Color-coded SPR sensograms demonstrate a concentration-dependent interaction between the wild-type C-terminal ABF FNBP4 fragment and immobilized G-actin, whereas (D) the mutant C-terminal ABF FNBP4 fragment exhibits markedly reduced binding to actin. For all measurements, association and dissociation phases were maintained at 120 s and 180 s, respectively, with buffer alone serving as the negative control. Sensogram data were globally fitted using a 1:1 Langmuir binding model, and the resulting binding responses for the wild-type and mutant C-terminal ABF FNBP4 fragments were plotted as response units (RU) versus protein concentration (µM).

To further corroborate the role of the KRRK motif (amino acids 1009-1012) within the C-terminal ABF FNBP4 (amino acids 946-1017) as the actin-binding site, surface plasmon resonance (SPR) binding analysis was performed using purified wild-type and mutant C-terminal ABF FNBP4 fragments. G-actin was first immobilized on the sensor chip surface (Fig. S 4), after which varying concentrations of the purified wild-type and mutant C-terminal ABF FNBP4 fragments were passed over the surface, with GST alone included as a negative control. GST exhibited no detectable response units across all tested concentrations (Fig. S 4), confirming the absence of nonspecific interaction with actin. Whereas, the resulting sensograms of wild and mutant C-terminal ABF FNBP4 revealed that, at equivalent protein concentrations, the wild-type fragment generated significantly higher response units than the mutant (Fig. 3C and D), indicating a substantial reduction in actin-binding affinity upon mutation of the KRRK motif. These SPR results are consistent with the TIRF-based actin polymerization assays, indicating binding to G-actin and inhibition of actin polymerization. Together, this strongly suggest that the KRRK motif within the C-terminal ABF region of FNBP4 is essential for both actin binding and functional inhibition of polymerization, underlining the regulatory significance of this region in actin dynamics.

### 2.5. FL-FNBP4 more potent inhibitor of FMN1

Previously, we reported that the N-terminal fragment of FNBP4 (WW1-WW2 FNBP4; 214-629 amino acids) inhibits FMN1-mediated actin cytoskeleton dynamics. Here, we show that the extreme C-terminal region (946-1017 amino acids) contains an actin-binding site and can suppress actin polymerization at higher concentrations. To investigate how this actin-binding activity contributes to FMN1 inhibition, we generated a full-length fragment, FL FNBP4 (214-1017 amino acids) (Fig. 4A), encompassing both the FMN1-interacting and actin-binding regions, and expressed and purified as a 6×His-tagged fusion protein from a bacterial system (Fig. 4C). We next assessed the effect of FL FNBP4 on actin polymerization using two FMN1 constructs, C-terminal FH1-FH2 FMN1, and C-terminal FH2 FMN1 (Fig. 4B and C). In bulk pyrene-actin assays, FL FNBP4 potently inhibited FH1-FH2 FMN1-mediated actin assembly in a concentration-dependent manner (Fig. 4D), with half-maximal inhibition of 50 nM FH1-FH2 FMN1 achieved approximately at 37 nM (Fig. 4E). To determine whether FL FNBP4 targets the conserved FH2 domain of FMN1, we tested the effects of FL FNBP4 on FH2 FMN1 activity (Fig. 4E). In contrast, FL FNBP4 inhibited FH2 FMN1 with reduced potency compared to FH1-FH2 FMN1 inhibition (Fig. 4G). This suggests that inhibition of FH2 FMN1 is substantially weaker, indicating a transient interaction with the FH2 domain in the absence of the FH1 domain and the interdomain linker region between the FH1 and FH2 domains of FMN1. Importantly, FL FNBP4 alone (200 nM) did not affect actin polymerization, confirming that its inhibitory activity is specific to FMN1-mediated assembly (Fig. 4F). Collectively, these findings establish that FL FNBP4 functions as a potent inhibitor of FMN1-driven actin polymerization, with preferential targeting of the FH1-FH2 construct. This inhibitory function is consistent with our previous observation that the WW1-WW2 FNBP4 fragment suppresses 50 nM FH1-FH2 FMN1 with half-maximal inhibition at 134 nM, whereas full-length FNBP4 exhibits greater potency, achieving half-maximal inhibition at 37 nM.

**Figure 4.**
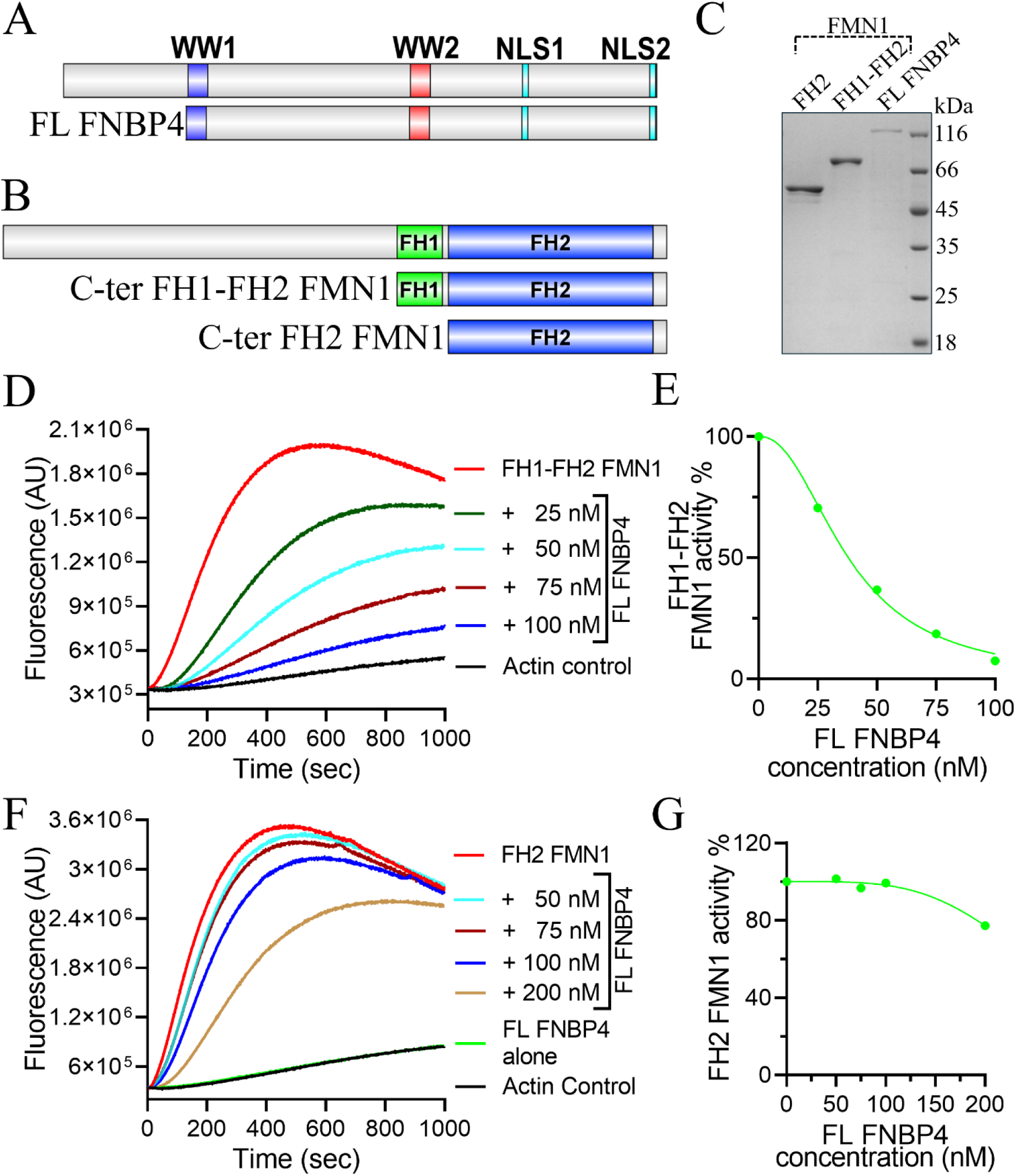
FL FNBP4 suppresses FMN1-driven actin polymerization in bulk assays. (A, B) Schematic representation of purified constructs used for in vitro experiments: FL FNBP4 (A) and FMN1 constructs, including C-terminal FH1-FH2 FMN1 and C-terminal FH2 FMN1 (B). (C) Coomassie-stained 10% SDS-PAGE showing purified C-terminal FH1-FH2 FMN1, C-terminal FH2 FMN1, and FL FNBP4 (residues 214–1017). (D, F) Pyrene-actin polymerization assays (2 µM actin, 10% pyrene-labeled) demonstrating the effects of FL FNBP4. Actin monomers were polymerized in the presence of 50 nM FH1-FH2 FMN1 with increasing concentrations of FL FNBP4 (D), or 50 nM FH2 FMN1 with increasing concentrations of FL FNBP4 (F). (E, G) Quantification of FL FNBP4-mediated inhibition under the conditions described in (D) and (F). Polymerization rates were determined from the slopes of the actin polymerization curves between 200 and 300 s. Percentage of FMN1 activity was calculated relative to the rate in the absence of FL FNBP4, and IC50 values were determined as the concentration of FL FNBP4 causing 50% inhibition of polymerization.

To further dissect the inhibitory effect of FL FNBP4 on FMN1-mediated actin polymerization, we performed spontaneous actin nucleation assays using TIRF microscopy. In these experiments, monomeric G-actin was incubated with either FH1-FH2 FMN1 or FH2 FMN1 in the presence or absence of FL FNBP4 (Fig. 5A). Both FMN1 constructs strongly promoted filament formation relative to actin alone (Fig. 5A and B). Expectedly in lower concentration of 200 nM FL FNBP4 alone did not alter filament numbers compared to the actin-only control (Fig. 5A). However, pre-incubation of FH1-FH2 FMN1 with FL FNBP4 prior to actin addition caused a substantial reduction in filament number relative to FH1-FH2 FMN1 alone (Fig. 5A and B). A similar, though less pronounced, inhibitory effect was observed when FH2 FMN1 was pre-incubated with FL FNBP4 (Fig. 5A and B). Moreover, we sought to further examine the effect of deleting the actin-binding region of full-length FNBP4 on FH1-FH2 FMN1-mediated actin polymerization. To this end, we generated a new fragment, ΔABF FL FNBP4 (214-1004 amino acids) (Fig. S 5A), which contains only the FMN1-interacting region and lacks the KRRK motif (1009-1012 amino acids). This construct was expressed and purified as a 6×His-tagged fusion protein from a bacterial expression system (Fig. S 5B). We next assessed the effect of 100 nM ΔABF FL-FNBP4 on 50 nM FH1-FH2 FMN1-mediated actin polymerization. We observed that the reduction in filament number was less pronounced than that observed with full-length FNBP4 (Fig. S 5C) and was comparable to the inhibition previously observed with the N-terminal WW1-WW2 FNBP4, due to steric hindrance. As expected, 100 nM ΔABF FL FNBP4 alone did not induce any significant change in filament number compared to the actin control (Fig. S 5C). These observations corroborate our bulk pyrene-actin assembly results and demonstrate that FL FNBP4 preferentially suppresses actin nucleation driven by the FH1-FH2 construct while exerting weaker inhibition on the isolated FH2 domain.

**Figure 5:**
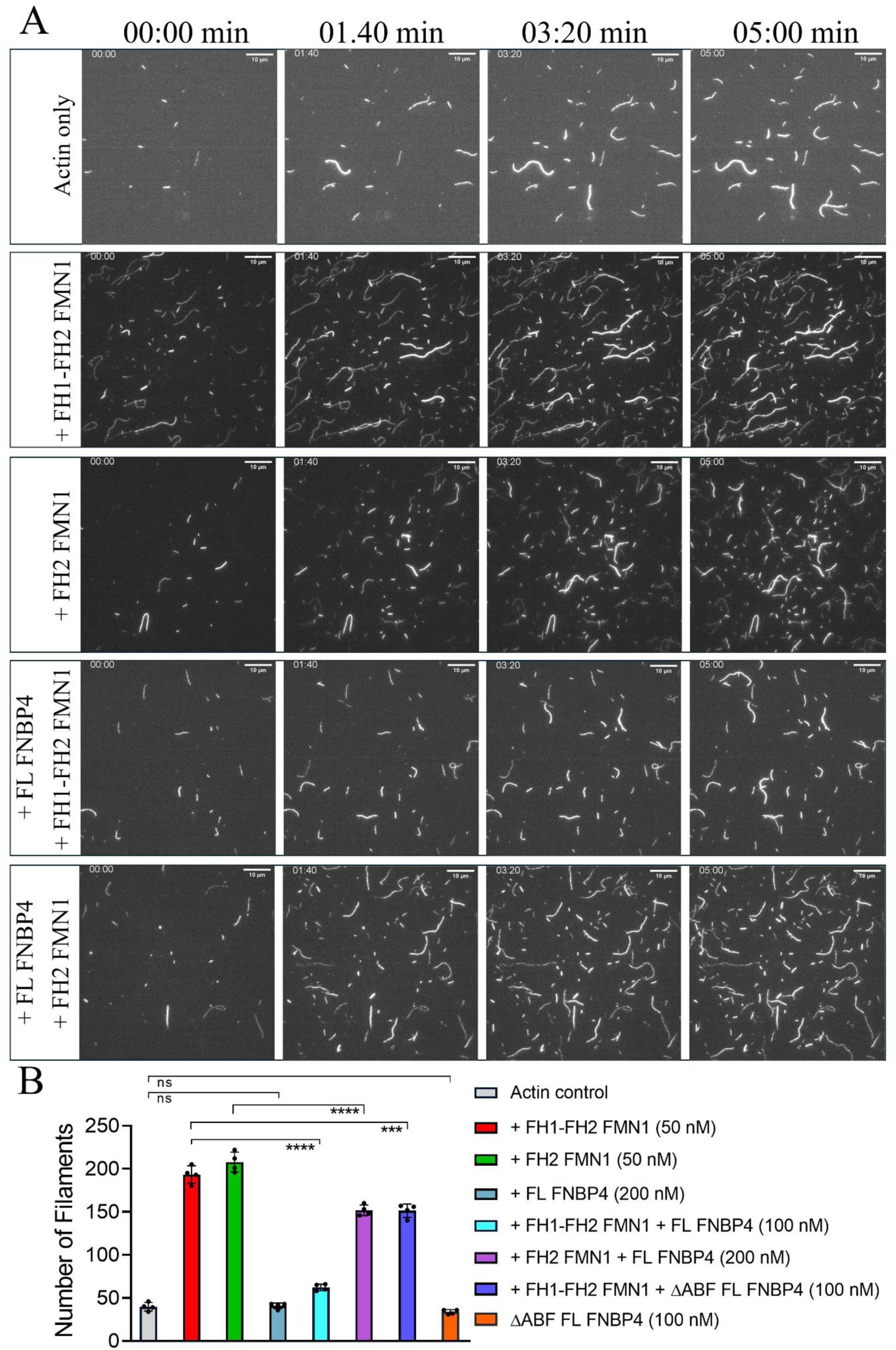
FL FNBP4 inhibits FMN1-mediated actin nucleation, observed by TIRF microscopy. (A) Time-lapse TIRF microscopy images showing actin filament assembly under five conditions: actin alone; 50 nM FH1-FH2 FMN1; 50 nM FH2 FMN1; 50 nM FH1-FH2 FMN1 with 100 nM FL FNBP4; and 50 nM FH2 FMN1 with 200 nM FL FNBP4. Scale bar, 10 µm. (B) Filament numbers at the endpoint were quantified from four independent fields. Error bars represent the standard deviation. Statistical analysis was performed using an unpaired two-tailed Student’s t-test (GraphPad Prism 8). Significance is indicated as *P ≤ 0.05, **P ≤ 0.01, ***P ≤ 0.001, ****P ≤ 0.0001, and ns (not significant).

### 2.6. FNBP4 Colocalizes with Actin in the Nuclear Compartment

Biochemical assays have demonstrated that FNBP4 directly interacts with both actin and FMN1. Our previous studies established that FNBP4 is predominantly a nuclear protein and identified two putative nuclear localization signals (NLSs), NLS1 (KGIKRKATEI; 787-797 amino acids) and NLS2 (RARLKRRKMA; 1005-1015 amino acids). Among these, only NLS1 was capable of directing nuclear localization, whereas NLS2 was insufficient. To further manifest the FNBP4 interaction with actin at the subcellular level. HeLa cells were transfected with a pmCherry-C1 actin-3×NLS P2A mCherry construct and immunostained for FNBP4 (Fig. 6A). Confocal imaging revealed substantial nuclear colocalization between FNBP4 and actin, with a Manders’ Colocalization Coefficient (M1) of 0.528 ± 0.08, supporting the notion that FNBP4 associates with actin within the nuclear compartment (Fig. 6A and B). These observations are consistent with our biochemical findings and suggest a potential functional role for the FNBP4 within the nucleus.

**Figure 6.**
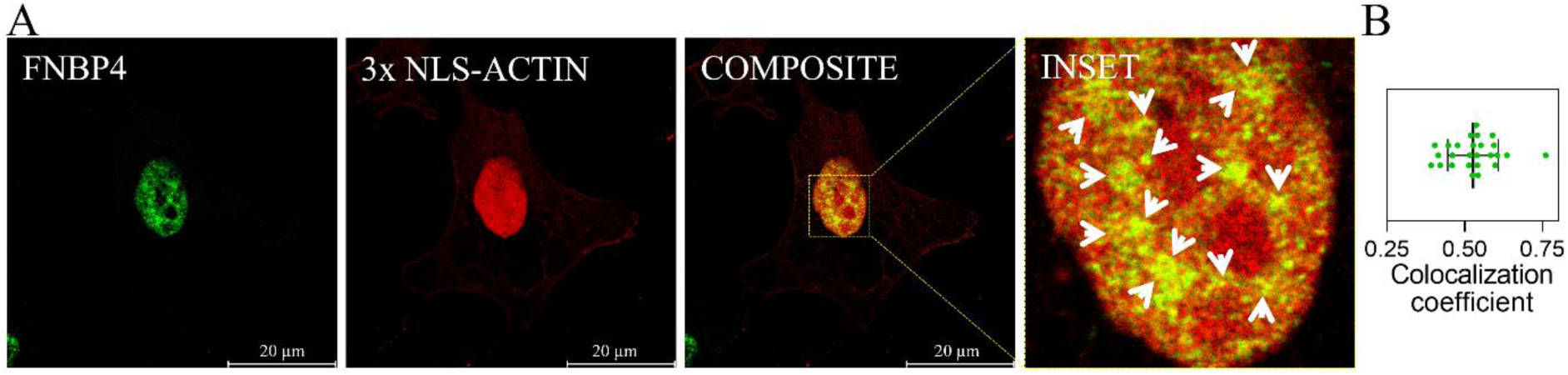
Nuclear colocalization of FNBP4 with actin in HeLa cells. (A) Cells were transfected with the pmCherry-C1 actin-3×NLS P2A mCherry construct. After transfection, the cells were fixed with 3.7% paraformaldehyde and subjected to immunofluorescence staining for FNBP4. FNBP4 was detected using an anti-FNBP4 antibody, and nuclei were counterstained with DAPI. The representative image shows a single optical section from the middle of the Z-stack. Scale bar: 20 µm. (B) Quantification of FNBP4 and actin colocalization using Manders’ Colocalization Coefficient. The scatter dot plot shows the distribution of data points along with standard deviations.

## 3. Discussion

In this study, we report for the first time that FNBP4 functions as an actin-binding protein. Specifically, its C-terminal region directly interacts with F-actin through a short basic motif (KRRK) located near the extreme C-terminus (1009-1012 amino acids) (Fig. 1). The actin-binding fragment of FNBP4 (C-ter ABF FNBP4; 946-1017 amino acids) exhibits a moderate affinity for F-actin (K_D_ ∼ 0.92 μM) (Fig. 1E). To further confirm the contribution of this motif, we compared the actin-binding ability of the wild-type C-ter ABF FNBP4 (KRRK) with that of the mutant form (AAAA) (Fig. 2). The mutant variant displayed a drastic loss of actin-binding activity, likely due to conformational changes caused by the replacement of basic residues. Moreover, the double-substitution variants, MutantA (KR-to-AA; 1009-1010 amino acids), MutantB (RR-to-AA; 1010-1011 amino acids), and MutantC (RK-to-AA; 1011-1012 amino acids), also lost their actin-binding ability (Fig. S3). These results indicate that disruption of any part of this short basic stretch is sufficient to abolish F-actin interaction.

This observation is consistent with previous studies demonstrating the significance of lysine-rich clusters in several actin-binding proteins. Notable examples include the KKEK motif in villin (Friederich et al., 1992), KKRKK or DAIKKK in cofilin (Iida et al., 1992; Yonezawa et al., 1989), the KSKLKKT motif in thymosin β4 (Safer et al., 1991), and the KKGGKKKG sequence in myosin (K, 1991). Mutational analyses of these motifs revealed that substitution of lysine residues significantly impairs actin binding. For instance, mutation of KKEK to KEEE in villin completely abolishes activity (Friederich et al., 1992), while substitution of KKRKK to KTLKK in cofilin markedly reduces actin-binding affinity (Iida et al., 1992). Consistent with this finding, our TIRF microscopy analysis revealed that the C-ter ABF FNBP4 inhibits actin polymerization, whereas the mutant fragment lacking the basic residues fails to exert this inhibitory effect (Fig. 3A and B). Complementing these functional observations, SPR binding analysis supports binding to G-actin and inhibition of actin polymerization, with mutation of the KRRK motif substantially reducing actin-binding affinity (Fig. 3C and D). Interestingly, in some cases, peptides or polymers containing similar basic amino acid stretches have been reported to promote G-actin to F-actin polymerization, likely through the formation of salt bridges between positively charged residues and actin monomers (Das et al., 2024a; Maiti et al., 2017). Collectively, our findings establish FNBP4 as a novel actin-binding protein and underscore that its C-terminal basic amino acid cluster serves as a conserved determinant for actin interaction, reinforcing the evolutionary importance of lysine-rich motifs in actin cytoskeleton regulation.

In our previous report, we demonstrated that FNBP4 localizes to the nucleus and identified two putative monopartite nuclear localization signals (NLS): KGIKRKATEI (787-797 amino acids; referred to as NLS1) and RARLKRRKMA (1005-1015 amino acids; referred to as NLS2) (Das et al., 2025). Functional dissection through overexpression of truncated GFP-tagged FNBP4 fragments (like NLS1 FNBP4; 1-1004 amino acids and NLS2 FNBP4; 801–1017 amino acids) revealed that NLS1 is responsible for the canonical nuclear localization of FNBP4 (Das et al., 2025). In contrast, NLS2 did not contribute to nuclear localization, suggesting it may serve a non-canonical function. Here, we established that the extreme C-terminal region of FNBP4 (1009-1012 amino acids; KRRK) is essential for its direct interaction with actin. Intriguingly, this actin-binding motif (KRRK) overlaps with the previously predicted non-canonical nuclear localization signal 2 (NLS2; 1005-1015 amino acids), which, although insufficient on its own for nuclear import, appears to play a critical role in mediating actin interaction within the nucleus. This is also supported by the overexpression of 3X-NLS actin and endogenous FNBP4 colocalization. These findings strengthen the idea of the potential role of NLS2 beyond nuclear import. Based on our observation that the GFP-tagged FNBP4 fragment encompassing amino acids 801-1017, which includes the NLS2 sequence, predominantly localizes to the cytoplasm. We can now assume that this is likely due to the interaction between the KRRK motif within NLS2 and cytosolic actin. It is plausible that actin binding masks the NLS2 motif, thereby interfering with its recognition by the nuclear import machinery. Given that proteins larger than ∼50 kDa typically require active transport through the nuclear pore complex (NPC), while smaller proteins can passively diffuse into the nucleus, the impaired nuclear translocation of the GFP-tagged NLS2 fragment (above 50 kDa) supports the hypothesis that actin binding occludes the NLS2 motif (Paine et al., 1975; Stewart, 2007). Taken together, these findings suggest that NLS2, despite containing a basic stretch of residues resembling a classical NLS, primarily function in actin binding. This adds an additional layer of complexity to the functional repertoire of FNBP4 and points to a potential role for actin interactions in modulating accessibility or functionality of nuclear targeting sequences in other proteins with similar motifs.

Nuclear resident proteins can generally be categorized into two groups. The first includes proteins that are constitutively translated in the cytoplasm and subsequently translocated into the nucleus, such as histones, CENP-A, and MYC (Bernardes and Chook, 2020; Chung and Levens, 2005; Quénet and Dalal, 2012). The second group consists of proteins whose nuclear localization is temporally regulated in response to specific stimuli, including JMY, FMN2, mDia2, FHOD1, and cofilin (Antoku et al., 2023; Belin et al., 2015; Iida et al., 1992; Ménard et al., 2006; Miki et al., 2009; Yamada et al., 2013; Zuchero et al., 2012). Both categories contain nuclear localization signals (NLSs), and in some cases, proteins harbor multiple NLSs, raising important questions regarding their functional relevance. Here, FNBP4 has two NLSs; their functions seem different, which emphasizes the functional disparity nature of NLS. Like cofilin, the KKRKK sequence serves both as an actin-binding site and an NLS during heat shock–induced nuclear accumulation (Iida et al., 1992). In contrast, the KRRK motif has been previously reported as a conserved NLS in various proteins, including Maf1, USP22, p100, and pUL48 (Brock et al., 2013; Liao and Sun, 2003; Pluta et al., 2001; Xiong et al., 2014). However, its role in nuclear import appears to be context-dependent. For instance, in the case of Retinitis Pigmentosa protein 2 (RP2), which contains multiple NLSs, the KRRK motif is not essential for nuclear localization (Hurd et al., 2011). These findings underscore the multifunctional nature of nuclear localization sequences, particularly in proteins that harbor more than one NLS. However, our findings align with previous observations from the Mullins lab and others, where several actin-binding proteins, such as JMY, ezrin, radixin, moesin, and talin, exhibit overlapping regions between their actin-binding site and NLS motifs (Batchelor et al., 2004; Borkúti et al., 2024; Fehon et al., 2010; Hemmings et al., 1996; Tidball and Spencer, 1993; Zuchero et al., 2012). This convergence suggests a broader regulatory principle wherein nuclear localization and actin binding are intricately linked through shared sequence elements. Further research is warranted to investigate the evolutionary and functional coupling of actin-binding site with nuclear localization signals, which may reveal new insights into the spatial regulation of cytoskeletal regulators and other dual-function proteins.

On the other hand, our findings establish a previously unrecognized dual functionality of FNBP4. We identified distinct roles for its terminal regions. Our earlier work showed that the N-terminal region interacts with FMN1 and inhibits FMN1-mediated actin remodeling (Das et al., 2025), while the newly identified C-terminal KRRK motif serves as an actin-binding site, revealing an unexpected additional layer of regulation. Intriguingly, we hypothesized that a fragment containing both functional sites would have a more pronounced effect on FMN1. Supporting this, we observed that FL FNBP4 strongly inhibits FH1-FH2 FMN1-mediated actin nucleation in vitro, with an IC50 value of 37 nM (Fig. 4D and E). Notably, this inhibitory effect is nearly fourfold stronger than that of the previously studied N-terminal (WW1-WW2) FNBP4 construct, which showed an IC50 of 134 nM. In addition, we also found that FL FNBP4 has some inhibitory effect on FH2 FMN1-mediated actin nucleation. However, FL FNBP4 strongly inhibits the FH1-FH2 FMN1 construct, while its impact on FH2 FMN1 is notably weaker, suggesting a functional disparity in its regulatory influence (Fig. 4 and 5). We speculate that this differential effect may arise from transient interactions in assay condition, as a similar weak inhibition was noted in our previous study (Das et al., 2025). However, we did not detect any direct biophysical interaction between FNBP4 and the FH2 domain of FMN1 by SPR analysis. Comparable multi-site inhibitory mechanisms have been reported for other formin regulators. For instance, Drebrin A inhibits mDia2 by interacting with both the FH2 domain and its tails (Ginosyan et al., 2019; Srapyan et al., 2023), while Hof1 suppresses Bnr1 activity through dual interactions between its F-BAR domain and the FH2 domain, as well as its SH3 domain and the FH1 domain (Garabedian et al., 2018; Graziano et al., 2014). Further structural studies will be necessary to elucidate the molecular basis of this regulatory phenomenon.

This dual functionality places FNBP4 among a growing group of formin-binding proteins such as Drebrin, Flii, Ena/VASP, Profilin, Spire, IQGAP1, and Anillin that regulate actin dynamics through the coordinated modulation of both filament assembly and formin activity. This dual capacity parallels other multifunctional actin regulators. For instance, Spire nucleates actin filaments via its WH2 domains while simultaneously cooperating with FMN-formins to coordinate filament elongation (Sitar et al., 2011; Vizcarra et al., 2011). Similarly, Ena/VASP proteins link actin monomers to filament barbed ends and engage formins through proline-rich motifs (Barzik et al., 2014, 2005; Breitsprecher et al., 2011). IQGAP1, on the other hand, anchors F-actin while modulating DIAPH1 activity (Chen et al., 2020; Liu et al., 2016; Mateer et al., 2004; Pimm et al., 2024). Unlike these activators, Drebrin A acts as an inhibitor by interacting with the FH2 domain and the C-terminal tail of mDia2, thereby suppressing its activity (Ginosyan et al., 2019). In addition, Drebrin A possesses actin-binding activity that contributes to its inhibitory role. Notably, Drebrin A is a neuron-specific protein that selectively inhibits diaphanous-related formins such as Daam1 and mDia2 but shows no effect on non-diaphanous formins (Ginosyan et al., 2019; Srapyan et al., 2023). In contrast, FNBP4 appears to have broader physiological significance as it is ubiquitously expressed and functions as an inhibitor of non-diaphanous formins. Further investigation is needed to determine whether FNBP4 can also inhibit diaphanous formins.

We observed that FNBP4 resides in and colocalizes with actin within the nucleus (Fig. 6). In line with this, previous studies have reported the nuclear presence of FMN1 (Chan and Leder, 1996; Isogai et al., 2015; Isogai and Innocenti, 2016). The nuclear localization of both FNBP4 and FMN1 raises intriguing questions about their regulation and dynamics. It remains to be determined whether these proteins shuttle between the nucleus and cytoplasm under specific physiological or stress conditions such as DNA damage, similar to JMY or FMN2 (Belin et al., 2015; Depraetere and Golstein, 1999; Yamada et al., 2013; Zuchero et al., 2012). This possibility is particularly compelling in light of earlier reports linking FNBP4 to the DNA damage response and its upregulation following γ-irradiation (Depraetere and Golstein, 1999). Functionally, our data reveal that FNBP4 predominantly acts as an inhibitor of the non-diaphanous formin FMN1, inhibiting its ability to drive actin nucleation, processive filament elongation, and filament bundling. This inhibitory function positions FNBP4 as a potential molecular brake that may play a crucial role in maintaining nuclear actin homeostasis. Based on our findings, we propose a refined version of the stationary inhibitory model previously described (Das et al., 2025). In this updated model, the N-terminal WW1 domain of FNBP4 interacts with the FH1 domain and the interdomain connector between the FH1 and FH2 domains of FMN1, thereby restricting the conformational flexibility required for efficient actin filament nucleation. Concurrently, the C-terminal region of FNBP4, which contains the KRRK motif, directly binds actin. We hypothesize that this dual interaction enables FNBP4 to transiently sequester actin monomers away from the FH2 domain of FMN1 during the early stages of filament formation. Such a mechanism would keep FMN1 in a restrained, stationary inhibitory state, allowing fine-tuned control over actin nucleation within the nucleus (Fig. 7).

**Figure 7.**
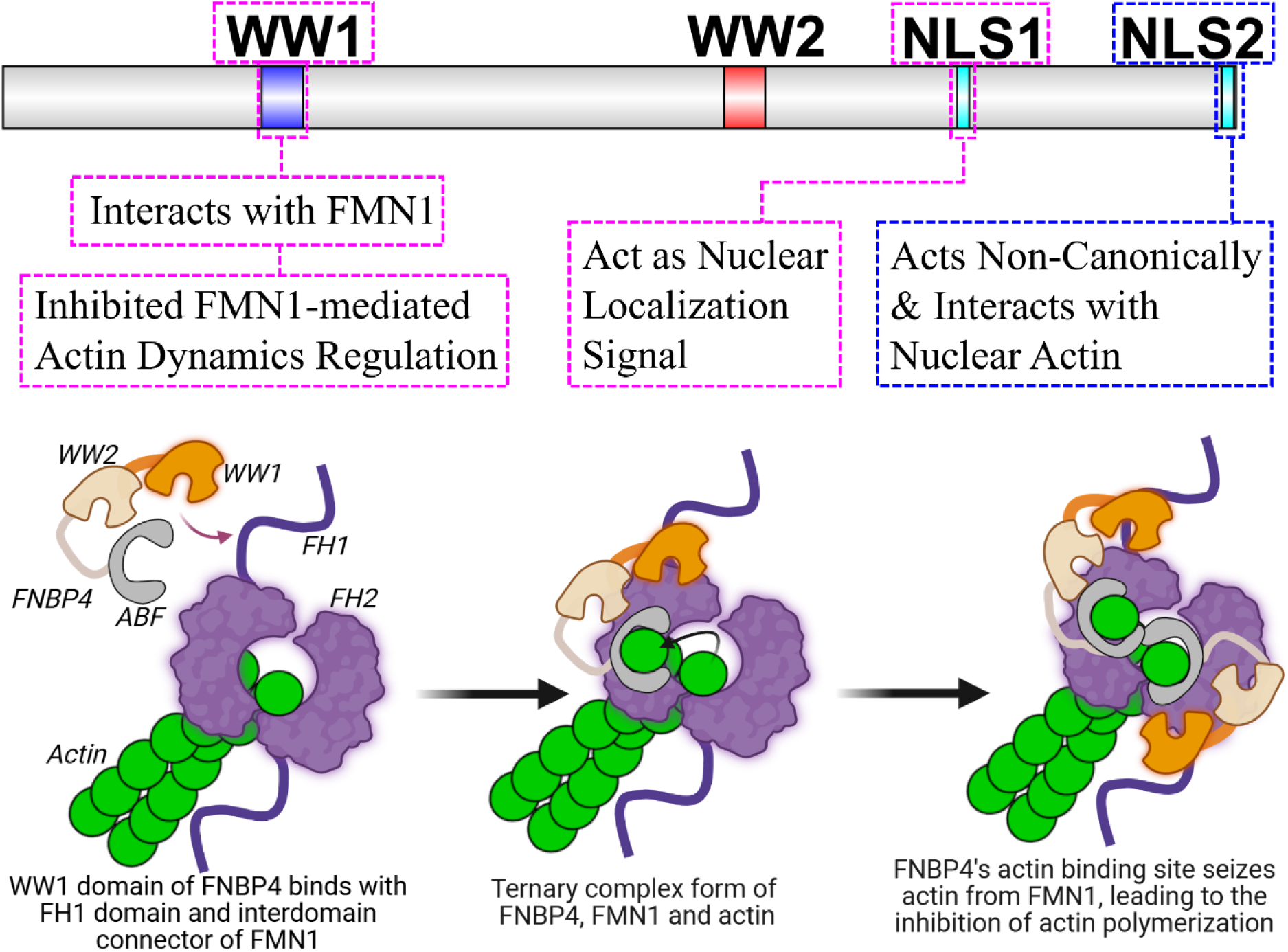
Proposed model illustrating FNBP4-mediated inhibition of FMN1-driven actin nucleation. The schematic represents our working model of how FNBP4 may regulate actin polymerization by interacting with FMN1. FMN1 is depicted in purple, with its FH1 domain and the interdomain connector between FH1 and FH2 shown as the primary interaction sites for FNBP4. The N-terminal WW1 domain of FNBP4 is shown in orange, engaging with these regions of FMN1. The C-terminal actin-binding fragment (ABF) of FNBP4 is illustrated in off-white, highlighting its direct interaction with actin, shown in green. We propose that this dual interaction allows FNBP4 to form a bridge between FMN1 and actin, potentially sequestering actin monomers away from the FH2 domain of FMN1 during the early stages of filament nucleation. This model provides a mechanistic basis for the observed inhibition of actin polymerization by the C-terminal ABF fragment of FNBP4.

Studying the nuclear actin cytoskeleton remains technically challenging, primarily due to the lack of nuclear-specific pharmacological probes. Despite these limitations, characterizing actin-binding proteins such as FNBP4 offers valuable insights into the regulation of nuclear actin dynamics. Further investigation into the interactions and regulatory mechanisms involving FNBP4 and nuclear actin nucleators such as FMN2, mDia2, FMN1, and FHOD1 could advance our understanding of nuclear actin-dependent processes. In conclusion, our study establishes FNBP4 as a previously unrecognized nuclear actin-binding protein that harbors a conserved basic motif critical for F-actin interaction, and functions alongside its formin-regulatory WW domain. This dual functionality opens new avenues for understanding the crosstalk between formin regulation and actin organization within the nucleus and underscores the need to explore nuclear-specific roles of formin-binding proteins in development and disease.

## 4. Materials and methods

### 4.1. Plasmid construct and cloning

Mouse FMN1 (plasmid pEGFPC2-FmnIso1a, Addgene #19320) and human FNBP4 cDNA (Dharmacon, MHS6278-202758566) were sourced for this study. Subsequently, different FMN1 and FNBP4 constructs were generated using methods described earlier (Das and Maiti, 2024; Das et al., 2025). Briefly, FMN1 truncation constructs comprising the C-terminal FH1-FH2 (870-1466 amino acids) and the C-terminal FH2 (983-1466 amino acids) were inserted into the pET28a(+) expression vector (Novagen). Moreover, FL-FNBP4 (214-1017 amino acids), ΔABF FL-FNBP4 (214-1004 amino acids), N-terminal FNBP4 (1-647 amino acids), C-terminal FNBP4 (647-1017 amino acids), and truncated C-terminal FNBP4 (732-904 amino acids) were cloned into the pET28a(+) vector, while the C-terminal ABF FNBP4 (946-1017 amino acids) was cloned into the pGEX-4T3 vector. For transfection studies, the pmCherry-C1 actin-3×NLS P2A mCherry plasmid was obtained from Addgene (RRID:Addgene_58475; deposited by Dyche Mullins).

### 4.2. Construction of Mutant Variants of the C-Terminal ABF FNBP4

Based on molecular docking predictions that identified the actin-binding interface at the extreme C-terminal region, we used standard PCR cloning to generate mutant variants by incorporating the desired base changes into the reverse primer. The mutants included C-terminal ABF Mutant FNBP4 (KRRK-to-AAAA; 1009-1012 amino acids), MutantA (KR-to-AA; 1009-1010 amino acids), MutantB (RR-to-AA; 1010-1011 amino acids), and and MutantC (RK-to-AA; 1011-1012 amino acids), all cloned into the pGEX-4T3 vector. Mutations were confirmed by DNA sequencing of the screened clones.

### 4.3. Protein purification

For 6x-His-tagged protein, Plasmid constructs encoding the C-terminal FH1–FH2 and C-terminal FH2 of FMN1, FL FNBP4, ΔABF FL-FNBP4 and N-terminal, C-terminal, and truncated C-terminal FNBP4 fragments were transformed into BL21 (DE3) E. coli (Agilent Technologies, #200131). Transformed cells were initially grown as primary cultures at 37°C in LB medium with 30 µg/mL kanamycin (Himedia). Secondary cultures were inoculated from these primary cultures and grown to an OD₆₀₀ of 0.5, followed by induction with 0.5 mM IPTG (Himedia). Expression of the FMN1 constructs and FL FNBP4 was carried out at 18°C for 12 h, whereas the N-terminal, C-terminal, and truncated C-terminal FNBP4 fragments were expressed at 25°C for 8 h. Cells were harvested by centrifugation and resuspended in lysis buffer (50 mM Tris, pH 8; 150 mM NaCl; 30 mM imidazole, pH 8; 0.5 mM DTT; 0.2% IGEPAL) supplemented with a protease inhibitor cocktail (aprotinin, pepstatin A, leupeptin, benzamidine hydrochloride, and PMSF). Cells were lysed on ice by sonication using an Ultrasonic Homogenizer (model U250, Takashi) at 1% amplitude with 10 s on/30 s off cycles for 4 min, and the lysate was clarified by centrifugation at 12,000 rpm for 10 min at 4°C to remove cell debris. The resulting supernatant was then incubated with Ni²⁺-NTA beads (Qiagen) for 2 h at 4°C on a ROTOSPIN™ rotary mixer at 5 rpm. Following incubation, the beads were washed three times with wash buffer (50 mM Tris, pH 8; 300 mM NaCl; 30 mM imidazole, pH 8) and bound proteins were eluted with elution buffer containing 50 mM Tris, pH 8; 20 mM NaCl; 350 mM imidazole, pH 8; and 5% glycerol. Finally, the eluted proteins were dialyzed for 4 h at 4°C against HEKG5 buffer (20 mM HEPES, 1 mM EGTA, 50 mM NaCl, and 5% glycerol).

For GST-tagged protein, Plasmid constructs encoding C-terminal ABF FNBP4, C-terminal Mutant ABF FNBP4, Mutant A, Mutant B, and Mutant C were transformed into BL21 (DE3) E. coli. Primary cultures of the transformed cells were grown at 37°C in LB medium containing 100 µg/mL ampicillin, and secondary cultures were inoculated from these primary cultures and grown to an OD₆₀₀ of 0.5, followed by induction with IPTG. Expression of all GST-tagged constructs was carried out at 25°C for 8 h. Cells were harvested by centrifugation and resuspended in lysis buffer (50 mM Tris, pH 7.5; 100 mM NaCl; 5 mM MgCl₂; 2 mM DTT; 0.2% IGEPAL; 5% glycerol; 1 mM EDTA) supplemented with a protease inhibitor cocktail. Cell lysis was performed on ice by sonication, and the lysate was clarified by centrifugation at 12,000 rpm for 10 min at 4°C. The resulting supernatant was incubated with glutathione agarose beads (GE Healthcare) for 2 h at 4°C under rotation at 5 rpm. After incubation, the beads were washed with wash buffer (50 mM Tris, pH 8; 150 mM NaCl; 1 mM EDTA) subsequently eluted with elution buffer (50 mM Tris, pH 8; 150 mM NaCl; 1 mM EDTA; 5% glycerol; 10 mM glutathione), and the bound proteins were subsequently dialyzed for 4 h at 4°C against dialysis buffer (20 mM HEPES, 1 mM EGTA, 50 mM KCl, and 5% glycerol).

### 4.4. Actin purification and labelling

Actin was purified from rabbit skeletal muscle powder following a published protocol (Das et al., 2025, 2024b; Dutta et al., 2026). Briefly, 1 g of acetone powder was suspended in 20 mL G-buffer (10 mM Tris-HCl pH 8, 0.2 mM CaCl₂, 0.2 mM ATP, 0.5 mM DTT, 0.1 mM NaN₃) and rotated at 4°C for 45 min. The mixture was centrifuged at 2000 rpm for 10 min, and the pellet was re-extracted once. Combined supernatants were filtered and polymerized with 2 mM MgCl₂ and 800 mM KCl, and F-actin was pelleted by ultracentrifugation at 65,000 rpm for 2h. The pellet was resuspended in G-buffer using dounce homogenizer, and dialyzed for 2 days at 4°C, cleared by ultra centrifugation at 80,000 rpm for 1 h, and purified by gel filtration on a HiPrep 16/60 Sephacryl S-200 HR column.

For Pyrene actin polymerization assay G-actin labelled with *N*-(1-pyrene)iodoacetamide. First G-actin was dialyzed against G-Buffer without DTT, then polymerized by adding 50 mM KCl and 2 mM MgCl₂, and subsequently incubated with a ten-fold molar excess of *N*-(1-pyrene)iodoacetamide targeting Cys-374 for 16 h at 4°C, as described previously (Doolittle et al., 2013; Kouyama and Mihashi, 1981).

### 4.5. Co-sedimentation assays

Actin (25 μM) was polymerized in F-buffer (10 mM Tris-Cl, pH 7.5, 0.2 mM MgCl₂, 0.2 mM CaCl₂, 50 mM KCl, 0.2 mM DTT, and 0.5 mM ATP) at 37 °C for 3 h and diluted to 5 μM in a 50 μL reaction volume. Test proteins (different FNBP4 constructs) were added at the indicated concentrations. Actin alone served as a control and sedimented into the pellet fraction, whereas protein in F-buffer without actin was used to assess solubility and remained in the supernatant fraction. After 15 min at room temperature, reactions were centrifuged at 90,000 rpm for 30 min at 4 °C (TLA100 rotor). Pellets were resuspended in 50 μL F-buffer, and equal volumes of pellet and supernatant fractions were mixed with sample loading buffer and analyzed by SDS-PAGE.

### 4.6. Binding affinity determination

The binding affinity of C-terminal ABF FNBP4 for actin was determined using a co-sedimentation assay with slight modifications from previously described protocols. Briefly, varying concentrations of C-terminal ABF FNBP4 (0.25, 0.5, 1, 2, 4, and 6 μM) were incubated with 2.5 μM actin. Following incubation, supernatant and pellet fractions were separated and run on separate gels simultaneously. Band intensities of the pelleted fractions were quantified using manually drawn rectangular regions of interest (ROIs) of equal size, large enough to encompass the largest band. Background signals were corrected by subtracting the average intensity of identically sized ROIs from three different regions of the gel. Corrected intensities were plotted against total C-terminal ABF FNBP4 concentration, and the resulting binding curve was fitted using a one-site specific binding model.

### 4.7. Molecular docking and Simulations

The full-length structure of FNBP4 (UniProt ID: Q8N3X1) was predicted using AlphaFold. The C-terminal actin-binding fragment (C-ter ABF FNBP4; 946-1017 amino acids) was extracted for docking studies. The structure of actin (PDB ID: 1J6Z) was obtained from the Protein Data Bank. The best model of FNBP4 was selected based on pLDDT and pTM scores and docked with actin using the HADDOCK server. The top-scoring docked complex was subjected to molecular dynamics (MD) simulations using GROMACS v2021. Hydrogen atoms were added, and the system was energy-minimized to remove steric clashes. The protein complex was placed in a cubic box and solvated with SPC216 water molecules. The system was neutralized with ions *via* the genion tool. Further system was equilibrated under constant temperature at 300 K, followed by a 50 ns production run.

Trajectory analyses were performed using GROMACS built-in tools. The root mean square deviation (RMSD) was calculated to assess structural stability, and the radius of gyration (Rg) was calculated to evaluate compactness. Plots were generated using GraphPad Prism v9.5. Protein-protein interactions were visualized using PyMOL v2.5, and 2D interaction maps were generated with LigPlot v2.2.

### 4.8. TIRF microscopy: Actin monomer sequestration and Actin nucleation

Glass coverslips washing and coating were performed as previously described (Das et al., 2024a; Mallick et al., 2025). Briefly, coverslips (22 × 22 mm) were cleaned with 2% Hellmanex III solution and coated with poly-L-lysine. Flow chambers were then assembled using double-sided tape on a glass slide, with the poly-L-lysine-coated coverslips facing inward. TIRF microscopy was performed on a Nikon Eclipse Ti2 with a 100× Apo TIRF oil objective, equipped with an ORCA-Flash4.0 Hamamatsu camera at room temperature.

For actin monomer sequestration assay, G-actin (3 μM) was converted from Ca²⁺-ATP-actin to Mg²⁺-ATP-actin in exchange buffer for 3 min. Actin was then incubated with C-terminal ABF FNBP4, C-terminal ABF mutant FNBP4 (4 μM each), or alone in KMEI buffer (50 mM KCl, 1 mM MgCl₂, 1 mM EGTA, 10 mM imidazole). Rhodamine-phalloidin was added for 30 s, and the reaction was subsequently diluted in TIRF buffer (10 mM imidazole, pH 7.4; 50 mM KCl; 1 mM MgCl₂; 1 mM EGTA; 0.2 mM ATP; 10 mM DTT; 1% methylcellulose; 10 μg/mL glucose oxidase; 20 μg/mL catalase; 15 mM glucose) for imaging at 5 and 10 min time points in separate experiments.

For actin nucleation assay, Mg²⁺-ATP-actin (2 μM) was mixed with either the preincubated FL FNBP4-FMN1 (FH1-FH2 FMN1 or FH2 FMN1) complexes, FL FNBP4 alone, or FMN1 (FH1-FH2 FMN1 or FH2 FMN1) alone in KMEI buffer. All other steps were the same, except images were acquired every 10 s for 10 min with the perfect focus system enabled during imaging.

### 4.9. Surface plasmon resonance

Interaction between G-actin, the C-terminal wild-type ABF FNBP4 (946-1017 amino acids), and mutant FNBP4 (KRRK-to-AAAA; 1009-1012 amino acids) were analysed using a BIAcore T200 system (GE Healthcare Life Sciences). Using the amine-coupling method, monomeric G-actin was immobilized on the surface of a CM5 sensor chip (Series S) (Wang et al., 2025). For immobilization, the sensor chip surface was first activated with an EDC/NHS mixture, followed by immobilization of 10 μg/mL G-actin in sodium acetate buffer (pH 4.5) in the presence of running buffer containing 5 mM HEPES (pH 7.4), 0.2 mM CaCl₂, and 0.2 mM ATP, to achieve a target response unit (RU) of 1000. After immobilization, the remaining reactive succinimide ester groups on the sensor surface were blocked using ethanolamine. Subsequently, different concentrations of purified C-terminal wild-type ABF FNBP4 (946-1017 amino acids) and mutant FNBP4 (KRRK-to-AAAA; 1009-1012 amino acids), along with buffer alone and/or GST only as a negative control, were injected over the G-actin-immobilized surface at a flow rate of 30 μL/min. The association and dissociation phases were set to 120 s and 180 s, respectively, and 10 mM glycine (pH 2.5) was flowed over the chip surface after each cycle for regeneration. HBS-EP buffer (10 mM HEPES, 30 mM EDTA, 150 mM NaCl, 0.05% surfactant P20, pH 7.4) was used as the running buffer during binding analysis. Binding data were analyzed using the resulting sensograms with BIAcore Evaluation Software (version 2.0). Kinetic parameters describing the interaction were obtained by applying global fitting of the sensogram data to a 1:1 Langmuir interaction model.

### 4.10. Pyrene actin polymerization assay

Actin polymerization was monitored using a fluorescence spectrometer (QM40, Photon Technology International, Lawrenceville, NJ) with 10% pyrene-labeled and 90% unlabeled monomeric actin in G-buffer. Reactions (2 µM actin) were converted to Mg²⁺-actin in exchange buffer (10 mM EGTA, 1 mM MgCl₂) and supplemented with FMN1 variants (C-terminal FH1-FH2 or FH2), FL FNBP4, or pre-formed FMN1–FL FNBP4 complexes at indicated concentrations (figure legends). Polymerization was initiated by 3 µL of 20× initiation mix in a final volume of 60 µL. Fluorescence was recorded at 25 °C (λ_ex = 365 nm, λ_em = 407 nm). The polymerization rate was derived from the slope between 200-300 s and used to determine IC₅₀ values.

### 4.11. Cell culture and transfection

HeLa cells (ATCC, catalog no. CCL-2; lot no. 70046455) were maintained in Minimum Essential Medium (MEM) supplemented with 10% fetal bovine serum, 2 mM L-glutamine, and 1% penicillin-streptomycin. For transfection, cells were seeded at a density of 5 × 10⁴ cells per well and transfected with the pmCherry-C1 actin-3×NLS P2A mCherry construct using Lipofectamine 2000 (Invitrogen) according to the manufacturer’s protocol. Transfected cells were incubated in transfection medium for 4 h, after which the medium was replaced with fresh complete medium and cells were cultured for an additional 18 h. Cells were then fixed with 3.7% paraformaldehyde for 15 min at room temperature, washed with PBS, and subjected to immunostaining for FNBP4.

### 4.12. Immunofluorescence staining and Imaging

For immunofluorescence staining of FNBP4, HeLa cells (pmCherry-C1 actin-3×NLS P2A mCherry construct-transfected) were fixed with 3.7% paraformaldehyde for 15 min at room temperature and washed with PBS. Then the cells were permeabilized with Triton X-100 (0.2 % in PBS) for 15 min at room temperature. Following permeabilization, cells were blocked with 2% BSA in PBS for 90 min at room temperature. After blocking, cells were incubated for 2 h at room temperature with mouse anti-FNBP4 (1:500) primary antibody, diluted in 1% BSA in PBST (PBS containing 0.075% Tween-20). After that, cells were incubated with the secondary antibody for 1 h at room temperature (Alexa Fluor™ 488–conjugated anti-mouse IgG, 1:1000), followed by nuclear staining with DAPI in Fluoroshield mounting medium, and coverslips were then mounted for imaging. Confocal images were acquired using a Leica SP8 microscope equipped with a 63×/1.40 N.A. oil immersion objective (HC PL APO CS2 63×/1.40 OIL). Colocalization was assessed using the JaCoP plugin in ImageJ (v1.54, https://imagej.net/ij/). Manders’ Colocalization coefficients were calculated from manually defined regions of interest (two per cell), using the thresholded fluorescence intensity for each channel.

### 4.13. Statistical analysis

Data were plotted and analyzed using GraphPad Prism (v8.4.2), with statistical significance determined by an unpaired, two-tailed Student’s *t*-test for two-group comparisons (*P* < 0.05; Fig. 3B and 5B).

## Data availability

All datasets that support the results of this study are available from the corresponding author upon reasonable request.

## Acknowledgements

This work was supported by the Anusandhan National Research Foundation (ANRF), DST, Government of India (Grant No. CRG/2023/006196 to S.M.). S.D. gratefully acknowledges the University Grants Commission for the fellowship. We thank the Department of Biological Sciences, IISER Kolkata, for access to the TIRF microscopy facility and computational resources. We also thank Amrita Maity and Saikat Das for assistance with molecular docking and simulation studies.

## Author contributions

**Shubham Das:** Data curation; investigation; visualization; validation; methodology; writing-original draft; writing-review and editing. **Saikat Das:** Data curation; investigation; visualization**. Sankar Maiti:** Conceptualization; funding acquisition; project administration; writing-original draft; writing-review and editing.

