## Supporting Information for "Dual Regulatory Role of Nuclear FNBP4 in Actin Binding and Formin FMN1 Inhibition"

**This PDF file includes**

Table S1 and S2

Figure S1 to S5

**Table S1: Detailed list of oligonucleotides used for cloning FMN1 and FNBP4 constructs.**

| Name of constructs | Vector | Primer sequences | Source |
|---|---|---|---|
| C-terminal FH1-FH2 FMN1 (870-1466 amino acids) | pET28a+ | FWD 5'-GGCGGATCCCCTCCAGCTCCCCCCACG-3'<br>REV 5'-GGACTCGAGTTAGTTGGTGGCCACGCTGGCT-3' | (Das and Maiti, 2024) |
| C-terminal FH2 FMN1 (983-1466 amino acids) | pET28a+ | FWD 5'-GGCGGATCCCGTAAACCAGCCATTGAGCCC-3'<br>REV 5'-GGACTCGAGTTAGTTGGTGGCCACGCTGGCT-3' | (Das and Maiti, 2024) |
| N-terminal FNBP4 (1-647 amino acids) | pET28a+ | FWD 5'-GGAGGATCCATGGGGAAGAAGTCCCGG-3'<br>REV 5'-GCCCTCGAGCTACTGTTTGGCAAGAGTCTCATCTC-3' | This paper |
| C-terminal FNBP4 (647-1017 amino acids) | pET28a+ | FWD 5'-GGAAGATCTTTGAAAGACAAAACTGGCACTG-3'<br>REV 5'-GGCCTCGAGCTATGTGTTTGGAGCCATTTTC-3' | This paper |
| Truncated C-terminal FNBP4 (732-904 amino acids) | pET28a+ | FWD 5' -GGCGGATCCGCGGAAGATGGTGAGATCC-3'<br>REV 5' -GGACTCGAGTTATTCTATAATGGTAGCGGTAGGC-3' | This paper |
| C-terminal ABF FNBP4 (946-1017 amino acids) | pGEX-4T3 | FWD **5'**-GGC**GGATCC**TTGGTAAAAAAGTGGCAGAGTATC-**3'**<br>REV 5'-GGCCTCGAGCTATGTGTTTGGAGCCATTTTC-3' | This paper |
| C-terminal ABF Mutant FNBP4 (KRRK-to-AAAA; 1009-1012 amino acids) | pGEX-4T3 | FWD **5'**-GGC**GGATCC**TTGGTAAAAAAGTGGCAGAGTATC-**3'**<br><br>REV 5'-GGACTCGAG**CTA**TGTGTTTGGAGCCAT<br>**TGCCGCTGCCGC**CAGCCTTGCTCTCCAATCCTCAGG-**3'** | This paper |
| MutantA (KR-to-AA; 1009-1010 amino acids) | pGEX-4T3 | FWD **5'**-GGC**GGATCC**TTGGTAAAAAAGTGGCAGAGTATC-**3'**<br><br>REV 5''-GGACTCGAG**CTA**TGTGTTTGGAGCCAT<br>**TTTCCT**TGCCGCCAGCCTTGCTCTCCAATCCTCAGG- 3'' | This paper |
| MutantB (RR-to-AA; 1010-1011 amino acids) | pGEX-4T3 | FWD **5'**-GGC**GGATCC**TTGGTAAAAAAGTGGCAGAGTATC-**3'**<br><br>REV 5''-GGACTCGAG**CTA**TGTGTTTGGAGCCAT<br>**TTTCGC**TGC**CTT**CAGCCTTGCTCTCCAATCCTCAGG-3'' | This paper |
| MutantC (RK-to-AA; 1011-1012 amino acids) | pGEX-4T3 | FWD **5'**-GGC**GGATCC**TTGGTAAAAAAGTGGCAGAGTATC-**3'**<br><br>REV 5''-GGACTCGAG**CTA**TGTGTTTGGAGCCAT<br>**TGCCGCTCTCTT**CAGCCTTGCTCTCCAATCCTCAGG- 3'' | This paper |
| FL-FNBP4 (214-1017 amino acids) | pET28a+ | FWD 5' -GGCGGATCCGGAATTGAGATGGGCGATTG-3'<br>**REV** 5'-GGCCTCGAGCTATGTGTTTGGAGCCATTTTC-3' | This paper |
| ΔABF FL FNBP4 (214-1004 amino acids) | pET28a+ | **FWD** 5' -GGCGGATCCGGAATTGAGATGGGCGATTG-3'<br>**REV** 5' – GGACTCGAG**TTA**CCAATCCTCAGGAAGGGCTTC-3' | This paper |

**Table S2: Detailed List of Antibodies and Recombinant DNA Constructs**

| Reagent and/or Resources | Reference or Source | Identifier or Catalog Number |
|---|---|---|
| **Antibodies** | | |
| Mouse Anti-FNBP4 | (Das et al., 2025) | N/A |
| Alexa Fluor™ 488-conjugated anti-mouse IgG | Invitrogen | A-11017 |
| **Recombinant DNA** | | |
| pET28+ | Novagen, Merck Life Science | 69864 |
| pGEX-4T3 | GE Healthcare Life Sciences | V010916 |
| pEGFPC2-FmnIso1a | Addgene | 19320 |
| Human FNBP4 cDNA | Dharmacon | MHS6278-202758566 |
| pmCherry-C1 actin-3×NLS P2A mCherry plasmid | Addgene | 58475 |

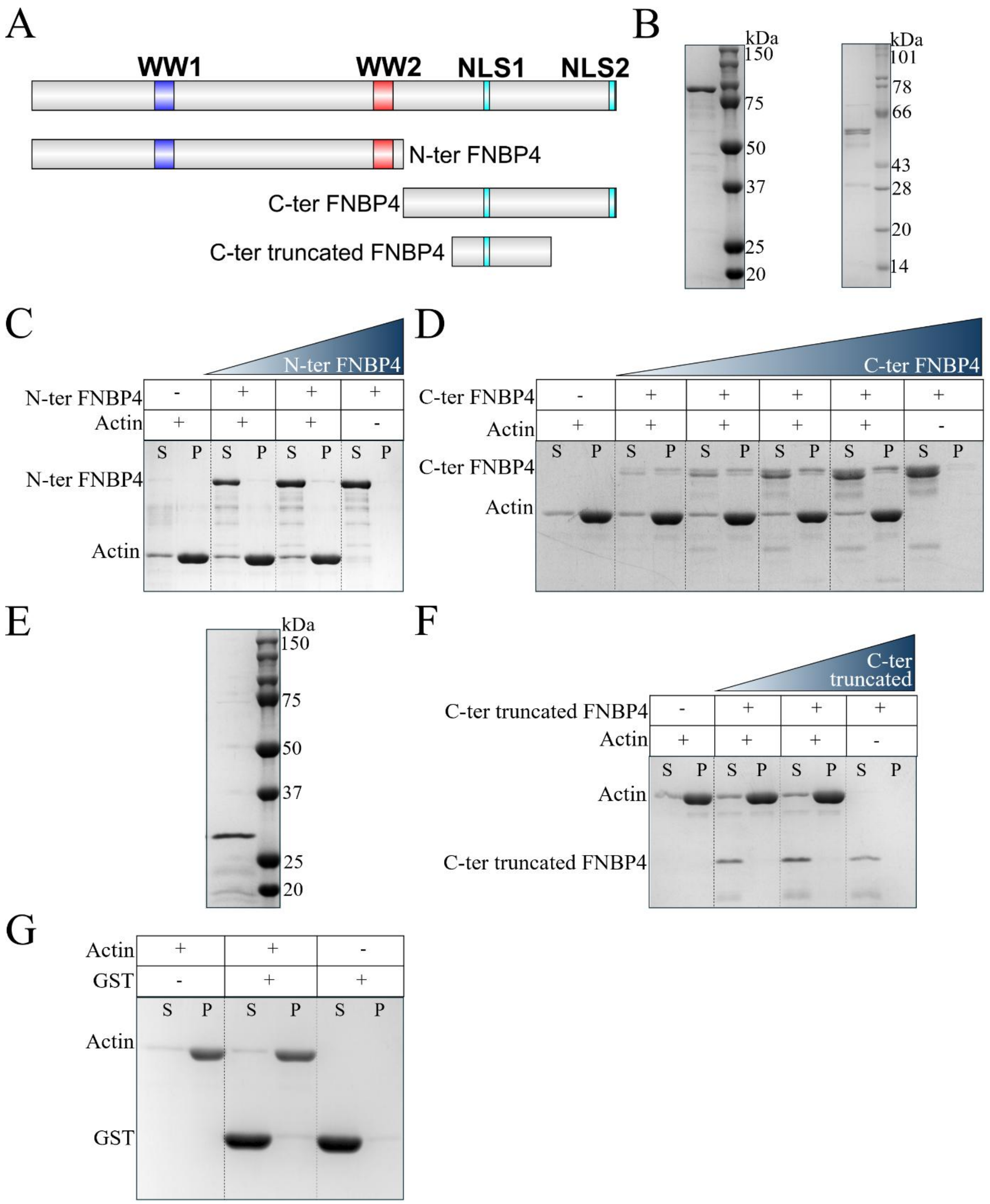


**Figure S1. Differential actin-binding properties of FNBP4 N- and C-terminal regions.** (A) Schematic of FNBP4 constructs used for in vitro studies, including N-terminal FNBP4 (residues 1-647 amino acids), C-terminal FNBP4 (residues 647–1017), and truncated C-terminal FNBP4 (residues 732-904 amino acids). (B) Coomassie-stained 10% SDS-PAGE showing purified N-terminal FNBP4 (1-647 amino acids) and C-terminal FNBP4 (647–1017) expressed as 6×His-tagged proteins. (C, D) F-actin co-sedimentation assays. Polymerized actin

(5 μM) was incubated in the absence or presence of N-terminal FNBP4 (1-647 amino acids) (C) or C-terminal FNBP4 (647-1017 amino acids) (D) for 15 min at room temperature. Samples were subjected to ultracentrifugation, and supernatant (S) and pellet (P) fractions were analyzed by 10% SDS-PAGE and visualized with Coomassie staining. N-terminal FNBP4 remained in the supernatant, showing no detectable binding to F-actin. In contrast, C-terminal FNBP4 was found in the pellet along with F-actin, indicating a specific interaction with actin. (E) Purification of truncated C-terminal FNBP4 (732–904 amino acids) shown by SDS-PAGE. F-actin co-sedimentation assay of truncated C-terminal FNBP4 (732–904 amino acids) (F) and GST (G). Both proteins remained in the supernatant after ultracentrifugation, indicating that C-terminal truncated FNBP4 and GST alone does not interact with F-actin.

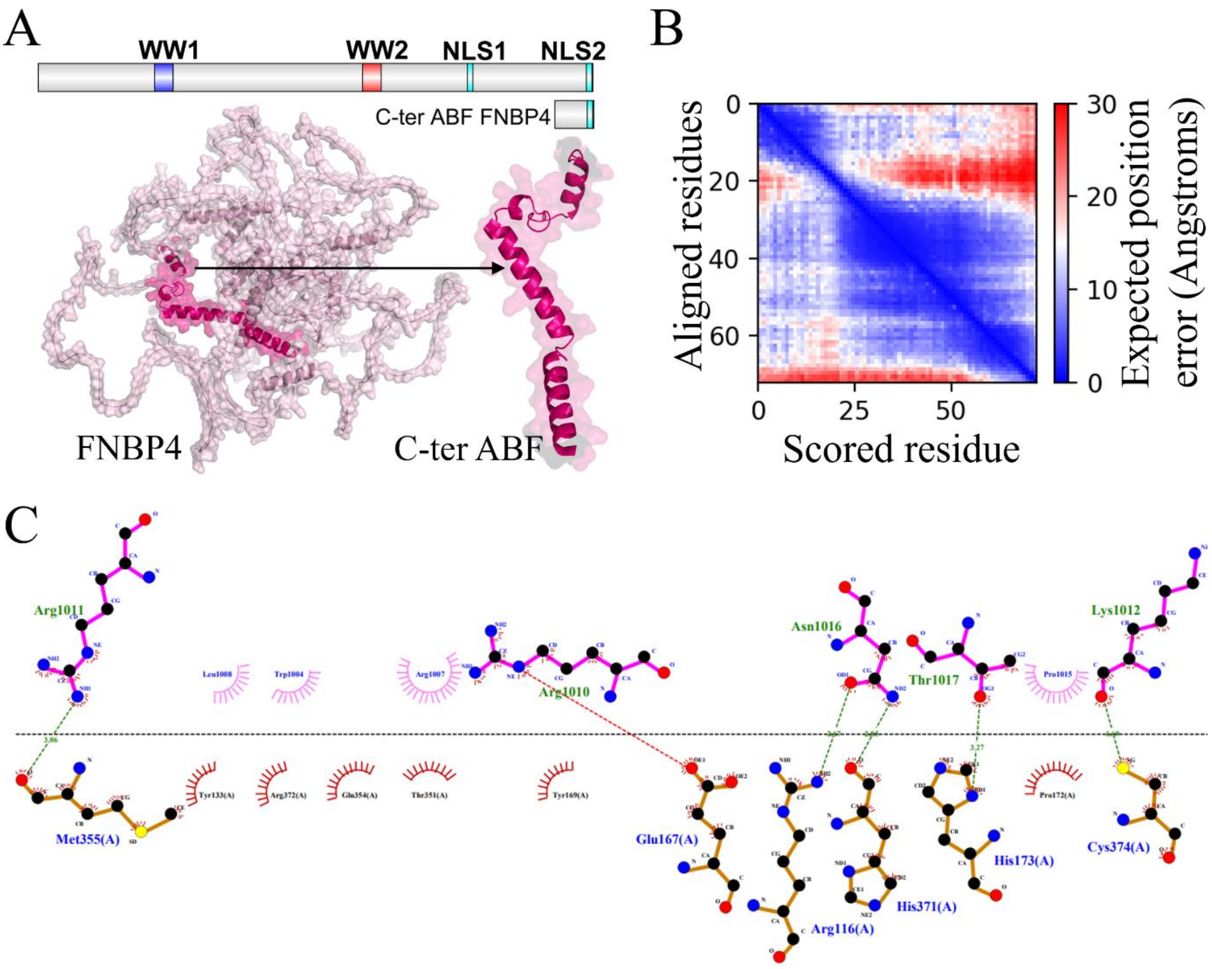


**Figure S2: Predicted 3D structure and interaction analysis of the C-terminal ABF region of FNBP4 with actin.** (A) AlphaFold-predicted 3D ribbon diagram of full-length FNBP4. The C-terminal ABF region of FNBP4 (946-1017 amino acids) used for docking is highlighted in light magenta, with three distinct helices. (B) Predicted Alignment Error (PAE) heatmap of the C-terminal ABF region (946-1017 amino acids) indicating model confidence. Blue regions correspond to high accuracy, while red regions indicate lower confidence. The predominantly blue regions align with the predicted α-helices, suggesting reliable structural modeling. (C) Interaction interface of the actin-C-terminal ABF FNBP4 complex analyzed using LigPlot+. Green dashed lines represent hydrogen bonds, and spokes indicate hydrophobic interactions between the two proteins.

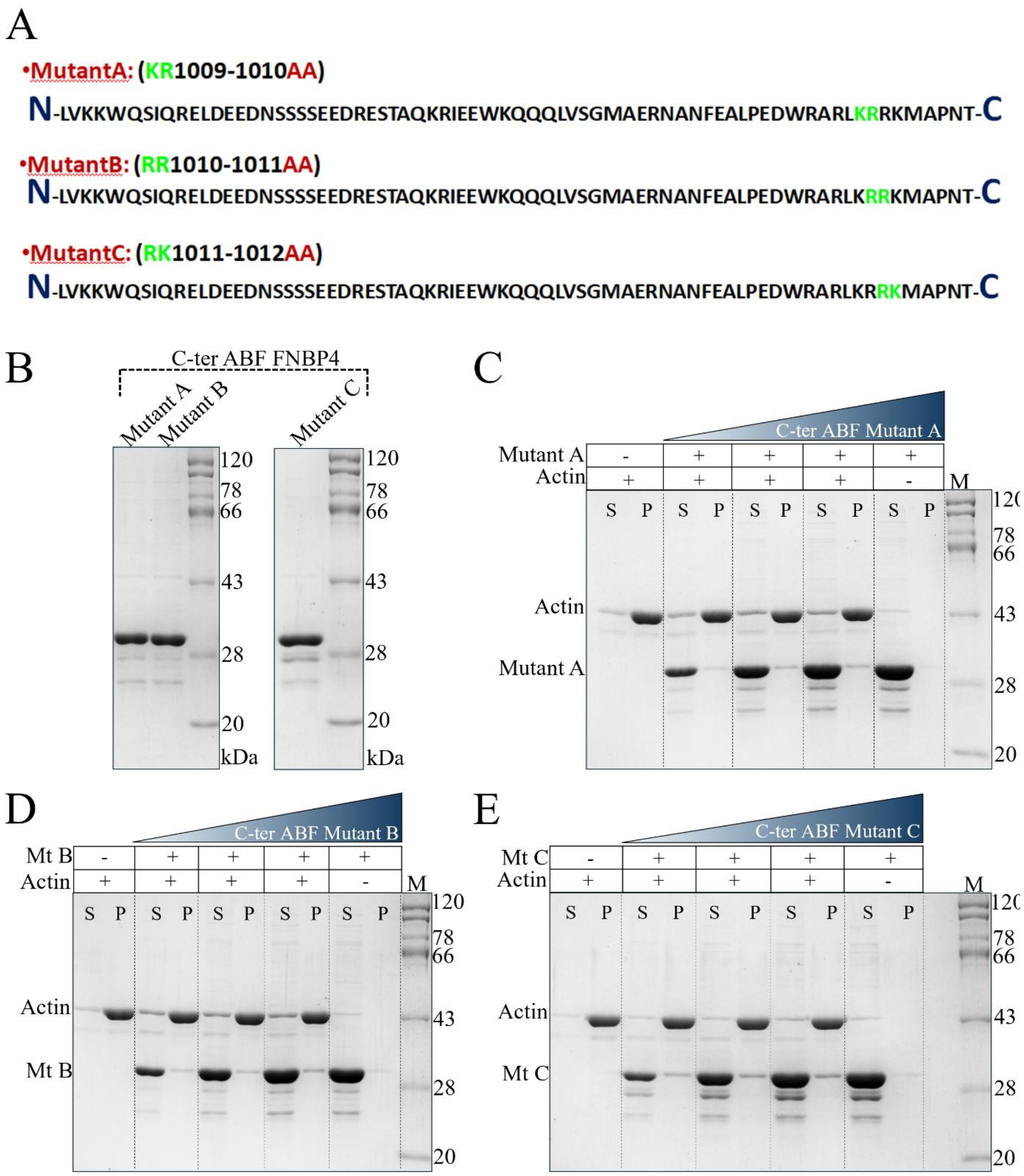


**Figure S3: The C-terminal KRRK motif is critical for FNBP4–actin interaction.** (A) Schematic of C-terminal ABF FNBP4 mutants highlighting substituted residues. MutantA (KR-to-AA; 1009-1010 amino acids), MutantB (RR-to-AA; 1010-1011 amino acids), and and MutantC (RK-to-AA; 1011-1012 amino acids) were generated to assess the actin binding activity. Mutated residues are shown in green, and alanine substitutions are highlighted in red. (B) 10% SDS-PAGE analysis of purified GST-tagged mutant proteins. (C-E) F-actin co-sedimentation assays for MutantA (C), MutantB (D), and MutantC (E). Polymerized actin (5 μM) was incubated with each mutant protein for 15 min at room temperature, followed by ultracentrifugation. Supernatant (S) and pellet (P) fractions were analyzed by SDS-PAGE and visualized by Coomassie staining. All mutants remained in the supernatant, indicating loss of actin-binding capability.

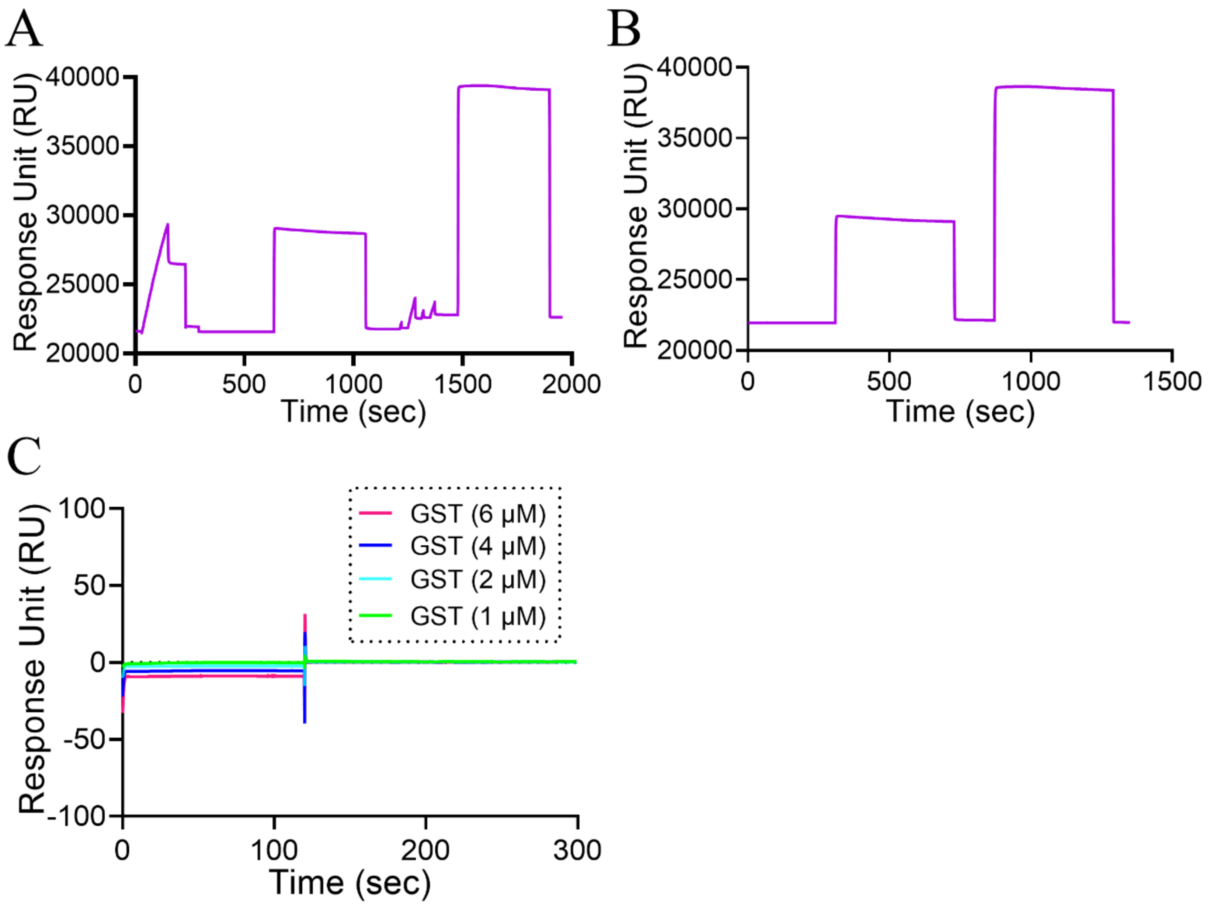


**Figure S4: Immobilization of G-actin and binding analysis demonstrates no detectable interaction with GST.** (A) Representative sensorgram showing immobilization of G-actin on reference channel 2 of the sensor chip. The surface was first activated using EDC/NHS, followed by injection of 10 µg/mL G-actin prepared in sodium acetate buffer (pH 4.5) for immobilization. Remaining active groups were subsequently blocked by ethanolamine. (B) Blank immobilization on reference channel 1, performed without G-actin, serving as a reference surface. (C) SPR sensorgrams from binding analysis using GST alone, demonstrating negligible response and confirming that GST does not interact with immobilized G-actin.

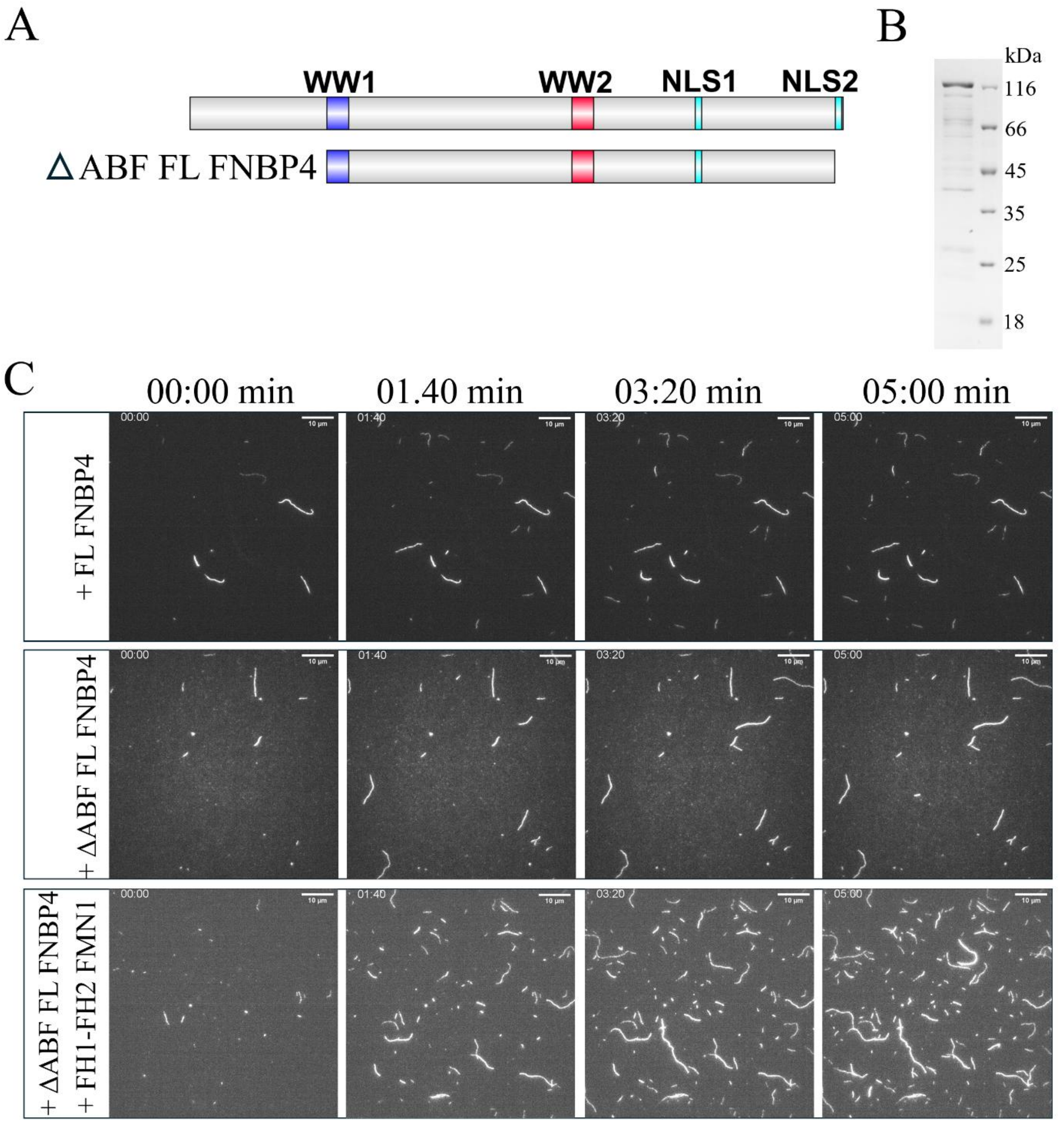


**Figure S5: ΔABF FL-FNBP4 (214-1004 amino acids) exhibits reduced inhibition of FMN1-mediated actin assembly.** (A) Schematic representation of the ΔABF FL FNBP4 (214-1004 amino acids) construct. (B) Coomassie-stained 10% SDS-PAGE gel showing purified ΔABF FL FNBP4 protein. (C) Time-lapse TIRF microscopy images showing actin filament assembly under three conditions: 200 FL FNBP4, 100 nM ΔABF FL-FNBP4 and 50 nM FH1-FH2 FMN1 with 100 nM ΔABF FL FNBP4. Scale bar, 10 µm.
